# The PopN gate-keeper complex acts on the ATPase PscN to regulate the T3SS secretion switch from early to middle substrates in *Pseudomonas aeruginosa*

**DOI:** 10.1101/2020.07.28.224923

**Authors:** Tuan-Dung Ngo, Caroline Perdu, Bakhos Jneid, Michel Ragno, Julia Novion Ducassou, Alexandra Kraut, Yohann Couté, Charles Stopford, Ina Attree, Arne Rietsch, Eric Faudry

## Abstract

*Pseudomonas aeruginosa* is an opportunistic bacterium of which the main virulence factor is the Type III Secretion System. The ATPase of this machinery, PscN (SctN), is thought to be localized at the base of the secretion apparatus and to participate in the recognition, chaperone dissociation and unfolding of exported T3SS proteins. In this work, a protein-protein interaction ELISA revealed the interaction of PscN with a wide range of exported T3SS proteins including the needle, translocator, gate-keeper and effector. These interactions were further confirmed by Microscale Thermophoresis that also indicated a preferential interaction of PscN with secreted proteins or protein-chaperone complex rather than with chaperones alone, in line with the release of the chaperones in the bacterial cytoplasm after the dissociation from their exported proteins. Moreover, we suggest a new role of the gate-keeper complex and the ATPase in the regulation of early substrates recognition by the T3SS. This finding sheds a new light on the mechanism of secretion switching from early to middle substrates in *P. aeruginosa*.

**Highlights:**

- T3SS substrates are secreted sequentially but information on the switches are missing
- Interaction of the T3SS ATPase with secreted proteins were investigated by different approaches
- Microscale Thermophoresis revealed a lower affinity for chaperones alone compared to complexes
- The Gate-keeper complex binds to the ATPase and increases its affinity for the needle complex
- A new role of the Gate-keeper complex is proposed, directly acting on the T3SS ATPase

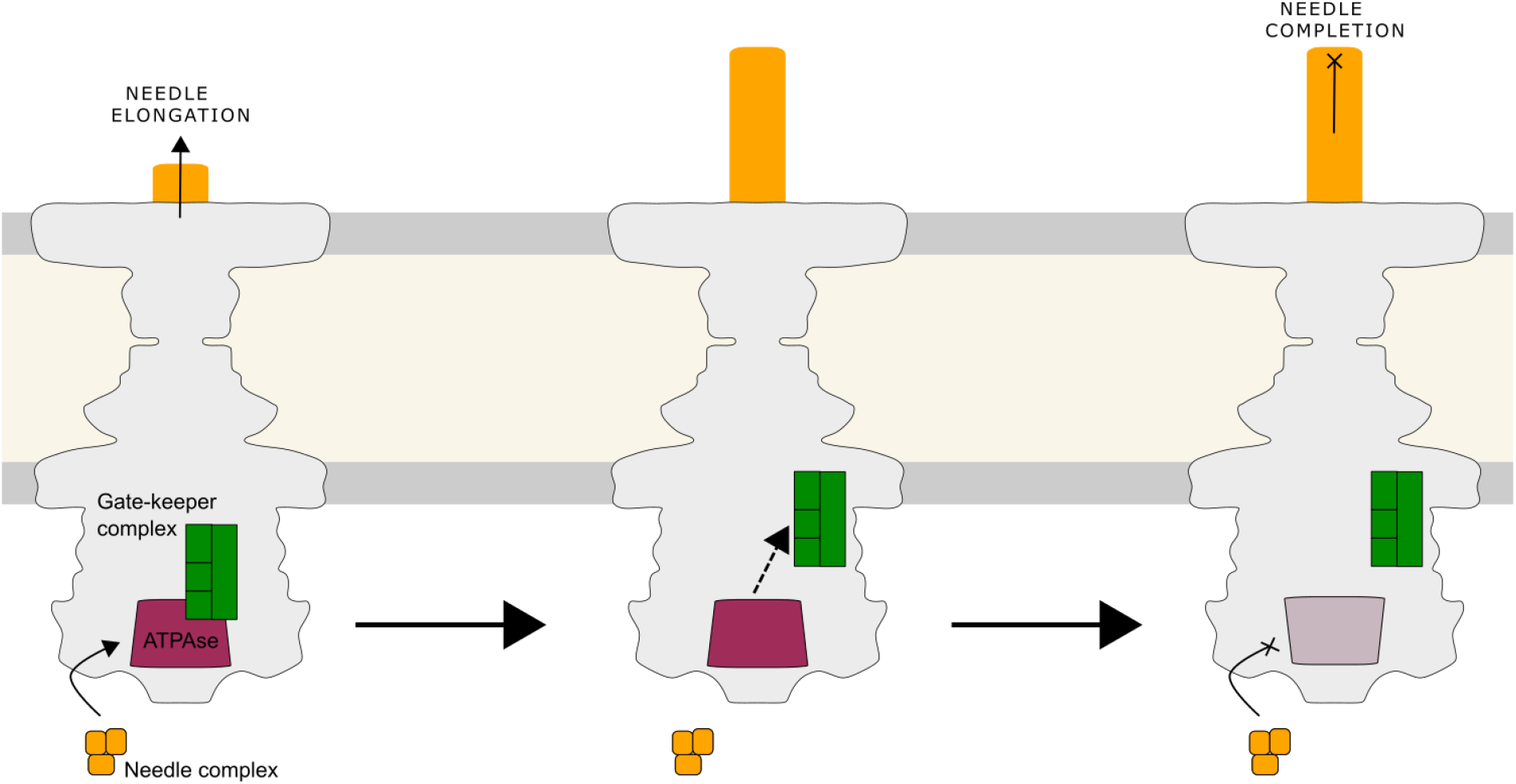

## Introduction

The Type III Secretion System (T3SS) is present in many Gram-negative bacteria, being responsible for the delivery of effector proteins directly into eukaryotic cells. An extracellular protrusion, the needle, connects the bacterial apparatus to the host membrane and cytoplasm. This nanomachine is constituted of more than 20 proteins mainly assembled in oligomers, which are located in the cytoplasm, inner membrane, periplasm, and outer membrane. To designate the T3SS components, the unified Sct nomenclature (Deng et al., 2017; Diepold and Wagner, 2014; Hueck, 1998) will be used throughout this work unless when describing our results or specific results from others. Table 1 describes the unified Sct names, the names in *P. aeruginosa*, the functional names of the principal proteins considered in this work and their counterparts in the closely-related flagellar apparatus.

**Table 1:** T3SS proteins identified by MS-based proteomics after pull-down with Strep-PscN.

| Unified Sct name | Name in *P. aeruginosa* | Functional name | Counterpart in the flagellar apparatus |
| --- | --- | --- | --- |
| SctN | PscN | ATPase | FliI |
| SctL | PscL | Peripheral stalk | FliH |
| SctO | PscO | Central stalk | FliJ |
| SctK | PscK | Accessory protein | - |
| SctQ | PscQ | Cytoplasmic ring | FliM–FliN |
| SctV | PcrD | Export gate | FlhA |
| SctF | PscF | Needle | FlgE |
| SctA | PcrV | Needle tip | - |
| SctB | PopD | Translocator | - |
| SctE | PopB | Translocator | - |
| SctU | PscU | Autoprotease switch protein | FlhB |
| SctP | PscP | Needle-length molecular ruler | FliK |
| SctW | PopN | Gate-keeper | - |

The conserved ATPase SctN is essential for T3SS function and bacteria deleted of the corresponding gene, *sctN*, display hampered T3SS secretion and cannot intoxicate eukaryotic cells. This enzyme is localized at the cytoplasmic extension of the T3SS and indirectly interacts with the membrane-associated proteins of the Type III C-ring SctQ and the basal body (Hu et al., 2015).

Displaying a structural homology to the F1-ATPases family, the SctN ATPase is believed to assemble into a hexameric ring presenting a central cavity that is blocked by the central stalk SctO which links the ATPase C-termini to the inner membrane export gate protein SctV (Gao et al., 2018; Hara et al., 2012; Ibuki et al., 2011, 2013; Lee et al., 2014; Majewski et al., 2019; Minamino et al., 2011). On the external side, the N-termini of SctN interact with the peripheral stalk SctL which acts as a stator and prevents the rotation of the ATPase (Halder et al., 2018; Imada et al., 2016).

The structure of the of SctN was characterized by X-ray crystallography and Cryo-EM, showing details of its three domains (Allison et al., 2014; Bernal et al., 2019; Burgess et al., 2016; Imada et al., 2007, 2016; Majewski et al., 2019; Zarivach et al., 2007). The C-terminal domain contains five α-helices and is thought to interact with the central stalk SctO and the secreted T3SS proteins in the first step of the secretion process. Indeed, specific single mutations in this domain abolish the binding of SctN to effector proteins (Allison et al., 2014). The central domain displays high similarity to the catalytic domain of other well-studied ATPases with a mixed α/β Rossmann fold displaying a parallel nine-stranded β-sheet flanked by three and four α-helices on either side. The Walker A and B motifs, which are crucial for the catalytic activity, are also found in this domain. Based on the similarity with the α/β subunit of the F1-ATPase, a model of the homo-hexameric ring SctN was built *in silico* with the ATP binding pocket interfaces located between two adjacent subunits (Allison et al., 2014; Bernal et al., 2019; Burgess et al., 2016; Demler et al., 2019). It was thus suggested that the oligomerization state is favorable to the catalytic activity. In addition to its role in the interaction with the peripheral stalk SctL, the N-terminal domain of SctN. was shown to be important for oligomerization and membrane association because the N-terminally truncated SctN is monomeric and catalytically less active (Burgess et al., 2016; Zarivach et al., 2007) and a single mutation in this domain impaired SctN ability to associate with the bacterial membrane (Akeda and Galán, 2004).

Functionally, T3SS ATPase has been proposed to be involved in the secretion process by acting in three different steps. First, it participates in the recognition of exported proteins at the entrance of the export apparatus. Secondly, upon hydrolyzing ATP, SctN has been shown to provide energy for the dissociation of chaperone-protein complexes, as shown for effector (Akeda and Galan, 2005; Lorenz and Buttner, 2009) and filamentous-tip proteins (Yoshida et al., 2014). Finally, energy from ATP hydrolysis is used to unfold secreted proteins in order to allow their passage through the narrow T3SS needle (Akeda and Galan, 2005). Besides ATP hydrolysis, the proton motive force (PMF) is thought to be the prominent energy source to push the export substrates through the needle through the interplay and complementarity between the ATPase and the export gate complex (Erhardt et al., 2014; Lee and Rietsch, 2015).

The interaction of SctN with effector proteins and their chaperones (Akeda and Galan, 2005; Akeda and Galán, 2004; Allison et al., 2014; Cooper et al., 2010; Gauthier, A. and Finlay, 2003; Lorenz and Buttner, 2009; Yoshida et al., 2014; Zarivach et al., 2007)), a tip-filamentous protein (Chen et al., 2013) and a translocon chaperone protein (Yoshida et al., 2014) was shown *in vitro*. Based on structural modelling, the chaperone of an effector was shown to dock to the loop of the two-helix-finger motif in the C-terminus of SctN (Allison et al., 2014). Thus, this region was pointed out as an important site for the recognition of secretion substrates by SctN. However, whether the exact binding site on SctN is identical for all exported proteins remains to be investigated.

In addition to these roles, it might be possible that SctN is involved in the regulation of the secretion process by participating in the sorting of substrates, thus allowing them to be hierarchically secreted. Indeed, during T3SS assembly, the needle subunits (early substrates) are first secreted, followed by the translocators (middle substrates) and then the effectors (late substrates) are secreted and translocated into host cells at the final stage of the secretion process. In the related flagellar machinery, the SctN-SctL (ATPase – Peripheral stalk) complex was shown to participate to the hierarchical sorting of the secreted protein (Inoue et al., 2018). Nonetheless, the sorting of substrates for secretion is a complex mechanism and undoubtedly requires an interplay between (the known, and maybe unknown,) regulator proteins such as SctP that controls the length of T3SS needle (Ho et al., 2017; Journet et al., 2003; Wagner et al., 2010), SctU that controls the switch from early to middle substrates (Edqvist et al., 2003; Monjarás Feria et al., 2015; Shen et al., 2012; Wood et al., 2008), SctW that controls the switch from middle to late substrates (Ferracci et al., 2005; Martinez-Argudo and Blocker, 2010; Roehrich et al., 2017; Yang et al., 2007), the needle tip protein SctA which may play the role of negative regulator of effector secretion in *Pseudomonas aeruginosa* and *Yersinia pestis* (Lee et al., 2014; Matson and Nilles, 2001; Nilles et al., 1997; Sundin et al., 2004) and other cytoplasmic proteins that were suggested as being the components of the substrate sorting platform SctO, SctK and SctQ (Lara-Tejero et al., 2011). Indeed, the global collaboration between these proteins and possibly the ATPase SctN to control the secretion process is still unclear.

In *P. aeruginosa*, the T3SS ATPase SctN is named PscN. In this study, full-length PscN was fused to tandem tags which promoted its high expression in a modified *P. aeruginosa* strain and allowed its purification. With an ELISA-based assay, we confirmed the interaction of PscN with a variety of secreted T3SS proteins and revealed for the first time the interaction with the SctF needle complex (PscE-PscF-PscG in *P. aeruginosa*). The dissociation constants of PscN and its partners were further determined by microscale thermophoresis (MST), showing a higher affinity between PscN and the protein complexes than with the chaperones alone. Finally, examining the relative affinities of PscN alone or bound to the SctW gate-keeper complex PopN-Pcr1-Pcr2-PscB, revealed that this complex can modulate partner recognition by PscN, suggesting a new mechanism by which PscN participates in the regulation of the secretion process.

## Results

### The ATPase PscN interacts *in vivo* with a wide range of T3SS proteins

The SctN ATPase is anchored to the T3SS apparatus through interactions with the central stalk SctO and the peripheral stalk SctL. To look for soluble partners of this protein in *P. aeruginosa*, full-length PscN was fused to a Strep-tag and expressed in the *P. aeruginosa* CHA strain harboring a chromosomal deletion of the *pscN* gene. After checking that the T3SS functionality was restored by this complementation (data not shown), Strep-PscN and its binding partners were enriched by affinity purification using streptactin beads and the corresponding fractions were analyzed by mass spectrometry (MS)-based proteomics. Among the identified proteins, several T3SS proteins were co-purified with PscN, including the exoenzymes S, T, Y, the translocators PopB (SctE), PopD (SctB) and their cognate chaperone PcrH, the peripheral stalk PscL (SctL), the export gate protein PcrD (SctV), the accessory protein PscK (SctK) and the chaperone PscB of the gate-keeper (SctW) complex (Table 2). In addition to structural partners, most of these partner proteins were shown to be secreted by the T3SS (Belyy et al., 2018; Goure et al., 2004; Maresso et al., 2006; Shen et al., 2008; Yahr et al., 1996; Yang et al., 2007) being consistent with a role of PscN in the recognition of exported protein as the first step of the secretion process.

**Table 2:** T3SS proteins identified by MS-based proteomics after pull-down with Strep-PscN.

| Protein name | Gene name | Peptides | Coverage | Spectral counts |
| --- | --- | --- | --- | --- |
| ATPase PscN (SctN) | *pscN* | 34 | 63.4% | 141 |
| Exoenzyme S (Exotoxin) | *exoS* | 7 | 19.7% | 9 |
| Peripheral stalk PscL (SctL) | *pscL* | 7 | 38.8% | 7 |
| Transcription activator ExsA | *exsA* | 6 | 14.0% | 6 |
| Transcription anti-activator ExsD | *exsD* | 5 | 20.3% | 5 |
| Translocator PopB (SctE) | *popB* | 4 | 12.3% | 4 |
| Adenylate cyclase ExoY (Exotoxin) | *exoY* | 3 | 10.3% | 4 |
| Translocator chaperone PcrH | *pcrH* | 3 | 20.4% | 3 |
| Major export apparatus protein PcrD (SctV) | *pcrD* | 2 | 3.5% | 3 |
| Exoenzyme T (Exotoxin) | *exoT* | 3 | 8.8% | 3 |
| Gate-keeper complex chaperone PscB | *pscB* | 1 | 7.1% | 1 |
| C ring protein PscQ (SctQ) | *pscQ* | 1 | 2.9% | 1 |
| transcription anti-antiactivator ExsC | *exsC* | 1 | 11.7% | 1 |
| Accessory cytosolic protein PscK (SctK) | *pscK* | 1 | 8.7% | 1 |
| Secretin PscC (SctC) | *pscC* | 1 | 2% | 1 |
| Translocator PopD (SctB) | *popD* | 1 | 3.4% | 1 |

### Fusion of a triple tag in a modified *P. aeruginosa* strain increased the yield of purified PscN under native conditions

The interaction of PscN with the peripheral stalk PscL was recently described using both proteins refolded from inclusion bodies (Halder et al., 2018). Using the construct employed for the pull-down study, several attempts failed to provide enough material for biochemical characterization. However, a fusion protein carrying one Flag and the Twin-Strep tag fused to the N-terminus of *pscN* (called FSS-PscN), was successfully expressed in *E. coli* BL21 and purified using Streptactin Sepharose (GE) beads. Strikingly, FSS-PscN showed a higher yield of purification than the Strep-PscN (Figure 1A). This could be due to the fact that the Twin-Strep tag increases binding to the affinity beads in comparison to a single Strep-tag. Moreover, the introduction of the triple tag lowers the pI of the protein from 7 to 6.2, possibly making it more soluble in the working buffer. Nonetheless, a high amount of one contaminant protein, identified as GroEL by MS-based proteomics (data not shown), was present above the band corresponding to PscN. The presence of GroEL suggested that the FSS-PscN folding is challenging in *E. coli*. We therefore tested the expression of FSS-PscN in *P. aeruginosa* ADD1976, which is an engineered *P. aeruginosa* strain adapted to the pET system (Table 4). Remarkably, a higher amount of purified protein and a lower contamination were observed in comparison to the expression in *E. coli* (figure 1A).

**Figure 1:**
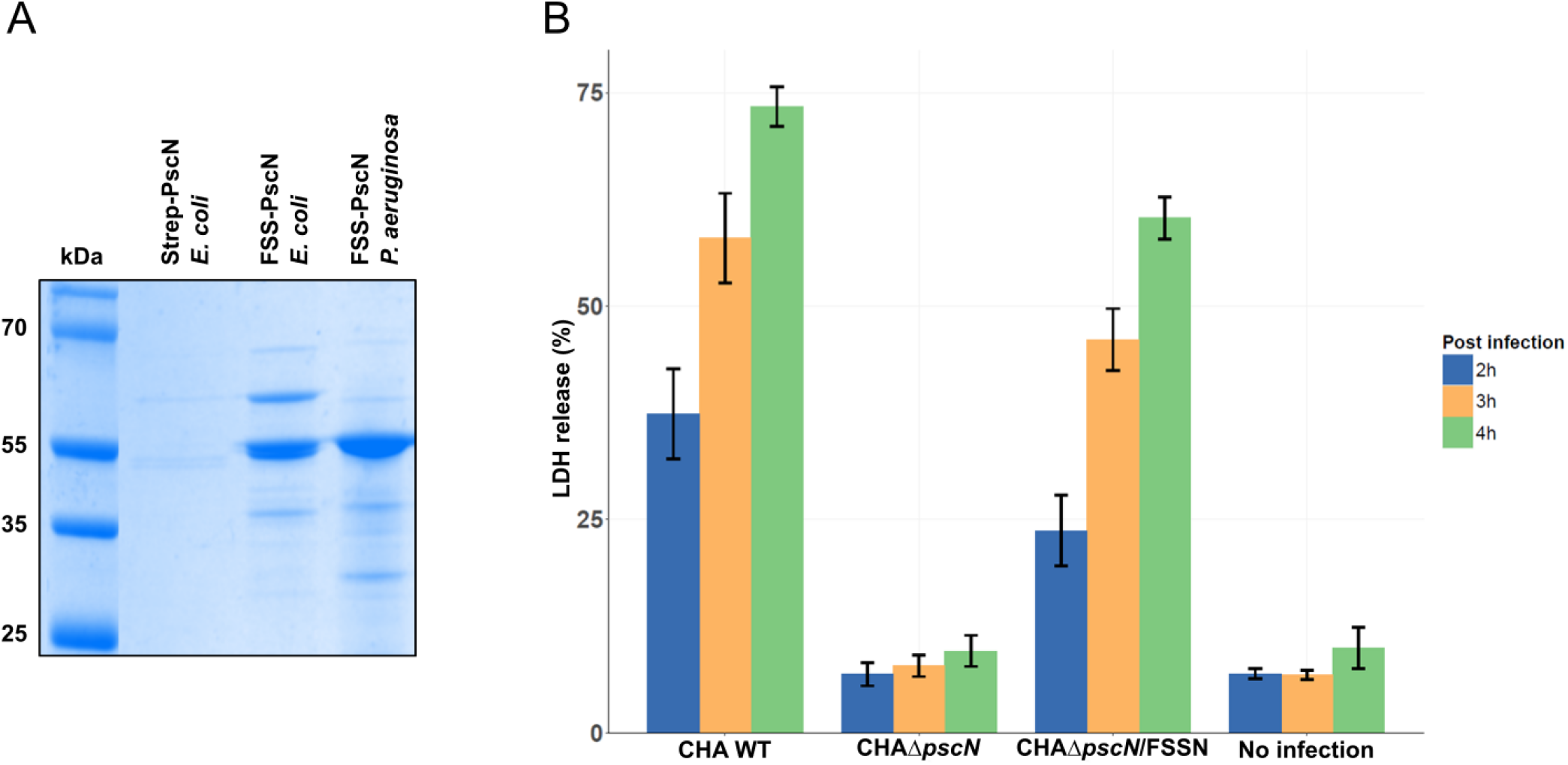
Fusion of a triple tag to PscN allows high purification yield upon expression in a modified P. aeruginosa strain and does not impair its activity. (A) Strep-PscN and FSS-PscN were expressed in E. coli BL21 and/or P. aeruginosa PAO1 ADD1976 and purified using Streptactin sepharose beads. Elution fractions from each condition were analyzed by Coomasie Blue. (B) Macrophages from the J774 cell line were incubated with P. aeruginosa CHA WT, ΔpscN or complemented strains at MOI of 10. Cell death was monitored at 2h, 3h and 4h post-infection by quantification of lactate dehydrogenase release. At each timepoint, the values obtained with the modified strains are statistically different from the ones obtained with the wt strain (ANOVA with Dunnet posthoc test, p-values < 0.05).

**Table 3:**
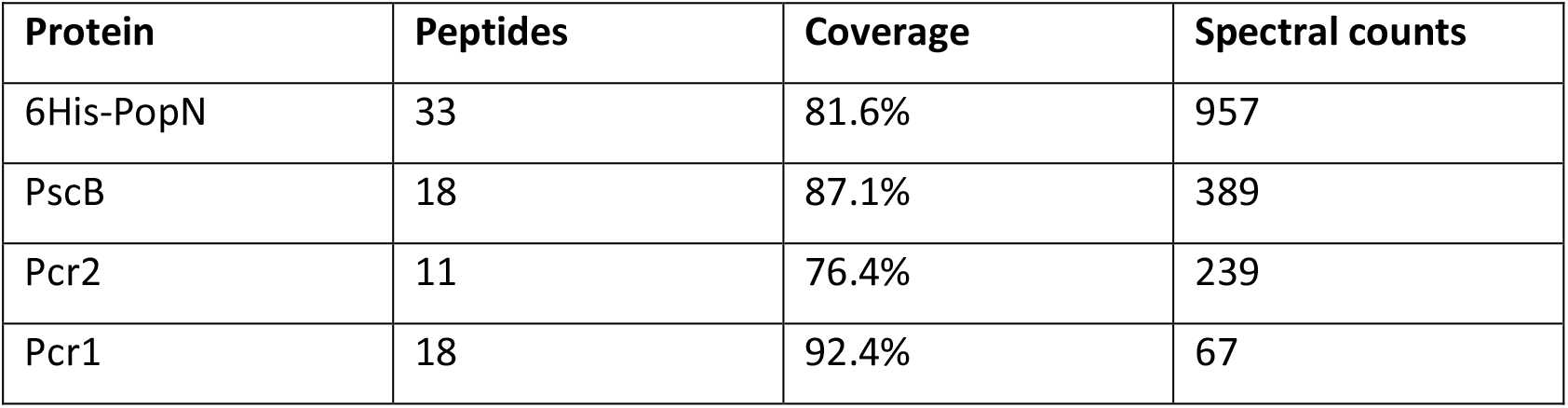
Characterization by MS-based proteomics of the recombinant PopN complex.

| Protein | Peptides | Coverage | Spectral counts |
| --- | --- | --- | --- |
| 6His-PopN | 33 | 81.6% | 957 |
| PscB | 18 | 87.1% | 389 |
| Pcr2 | 11 | 76.4% | 239 |
| Pcr1 | 18 | 92.4% | 67 |

**Table 4:** Strains and plasmids used in this work.

| **Strain** | **Description** | **Source or reference** |
| --- | --- | --- |
| **CHA $\Delta pscN$** | CHA *pscN* deletion mutant | This study |
| **CHA $\Delta pscN$/pIApG-Strep-PscN** | $\Delta pscN$ carrying Strep-PscN in pIApG | This study |
| **CHA $\Delta pscN$/pIApG-Flag-Strep-Strep-PscN** | $\Delta pscN$ carrying Flag-Strep-Strep-PscN in pIApG | This study |
| **PAO1 ADD1976/pET15bVP-Flag-Strep-Strep-PscN** | Flag-Strep-Strep-PscN expressed in modified PAO1 strain and in modified pET15b (N-terminal FSS) | This study (Arora et al., 1997; Brunschwig and Darzins, 1992) |
| **BL21 (DE3)/pET15b-Flag-Strep-Strep-PscN** | Flag-Strep-Strep-PscN expressed in BL21 and in pET15b (N-terminal FSS) | This study |
| **BL21 (DE3)/pET15b-Strep-PscN** | His-Strep-PscN expressed in BL21 and in pET15b (N-terminal His Strep) | This study |
| **BL21 (DE3)/pET15b-PscE-PscF-PscG** | PscE-PscF-PscG complex expressed in BL21 and in pET15b (N-terminal His) | (Quinaud et al., 2005) |
| **BL21 (DE3)/pET15b-PscE-PscG** | PscE- PscG complex expressed in BL21 and in pET15b (N-terminal His) | (Quinaud et al., 2005) |
| **BL21 (DE3)/pET15b-ExoU** | ExoU expressed in BL21 and in pET15b (N-terminal His) | (Gendrin et al., 2012) |
| **BL21 (DE3)/pET15b-SpcU** | SpcU expressed in BL21 and in pET15b (N-terminal His) | (Gendrin et al., 2012) |
| **BL21 (DE3)/pET Duet-ExoU-SpcU** | ExoU-SpcU complex expressed in BL21 and in pET15b (N-terminal His) | (Gendrin et al., 2012) |
| **BL21 (DE3)/pET15b-PcrV** | PcrV expressed in BL21 and in pET15b (N-terminal His) | (Goure et al., 2004) |
| **BL21 (DE3)/pET15b-PcrH** | PcrH expressed in BL21 and in pET15b (N-terminal His) | (Schoehn et al., 2003) |
| **BL21 (DE3)/pET30b-PcrH-PopB** | PcrH-PopB complex expressed in BL21 and in pET15b (N-terminal His) | (Schoehn et al., 2003) |
| **BL21 (DE3)/pET30b-PcrH-PopD** | PcrH-PopD complex expressed in BL21 and in pET15b (N-terminal His) | (Schoehn et al., 2003) |
| **BL21 codon plus “RP”/pET28bTEV-His-PopN-Pcr1-Pcr2-PscB** | PopN complex expressed in BL21 and in pET28b (N-terminal His) | This study |

To ascertain whether the relatively long tag (50 residues) could affect the conformation and function of the protein, we compared the cytotoxicity of the wild-type strain to the one of a deletion mutant strain *ΔpscN* complemented with *fss-pscN* expressed under the control of a T3SS promoter. For this purpose, the cytotoxicity of the different strains was tested on the J774 macrophage cell-line through measurement of the release of lactate dehydrogenase (LDH) to monitor cell death. Indeed, the complementation with FSS-PscN almost fully restored the virulence potential of *P. aeruginosa* mutant strain *ΔpscN* (Figure 1B), thus indicating that the introduction of one Flag and two Strep tags is not deleterious to the activity of PscN.

Furthermore, FSS-PscN displayed a Mg^2+^ or Zn^2+^-dependent ATPase activity that peaked at pH 9 (Figure S1), confirming that the full-length purified FSS-PscN was catalytically active.

### PscN interacts with effector, translocator, needle and gate-keeper proteins

PscN was shown to interact with the peripheral stalk PscL (SctL) (Halder et al., 2018) while SctN from other species was shown to interact with several classes of exported substrates, such as effector, translocator and gate-keeper proteins (Akeda and Galan, 2005; Akeda and Galán, 2004; Allison et al., 2014; Botteaux et al., 2009; Cooper et al., 2010; Gauthier, A. and Finlay, 2003; Lorenz and Buttner, 2009; Zarivach et al., 2007). To further explore the binding spectrum and specificities of PscN, an ELISA-based assay was performed with purified FSS-PscN and purified partner proteins including effector, translocator, needle and gate-keeper proteins (the cargos) and their chaperones in complex or alone.

Four classes of T3SS complexes were selected: i) the effector ExoU and its chaperone SpcU whose crystallographic structure was solved (Gendrin et al., 2012), ii) the translocators PopB and PopD and their cognate chaperone PcrH that have been extensively characterized (Discola et al., 2014; Faudry et al., 2007; Goure et al., 2004; Job et al., 2010); iii) the PscF (SctF) needle subunit that associates with the co-chaperones PscE and PscG (Ple et al., 2010; Quinaud et al., 2005, 2007); iv) The gate-keeper complex is composed of the gate-keeper PopN (SctW) associated with the co-regulator Pcr1 (TyeA in *Yersinia* spp.) and the two chaperones Pcr2 and PscB (Schubot et al., 2005; Yang et al., 2007). MS-based proteomic characterization of the proteins co-purified with 6His-tagged PopN indicated that it was effectively associated with its three partners (Table 3). Indeed, the interactions within this complex were mapped *in vivo* using a bacterial two-hybrid system showing that Pcr1 interacts with PopN, Pcr2 with PscB and that Pcr2 and PscB are needed to interact with PopN (Figure S2).

The potential partner proteins fused to a 6His-tag at their N-terminus were purified on a Nickel affinity column, followed by size exclusion chromatography (SEC). These proteins were then coated onto an ELISA plate and Bovine Serum Albumin (BSA) and PBS were used as negative controls. Subsequently, FSS-PscN was incubated at four different concentrations and its binding was detected by anti-PscN and secondary HRP-conjugated antibodies. The signal intensity was much higher with the T3SS proteins in comparison to negative controls and was dependent on the concentration of PscN (Figure 2). These results indicated that PscN interacts with the tested proteins, thus confirming its role in the recognition of exported proteins for the first step of the secretion process.

**Figure 2:**
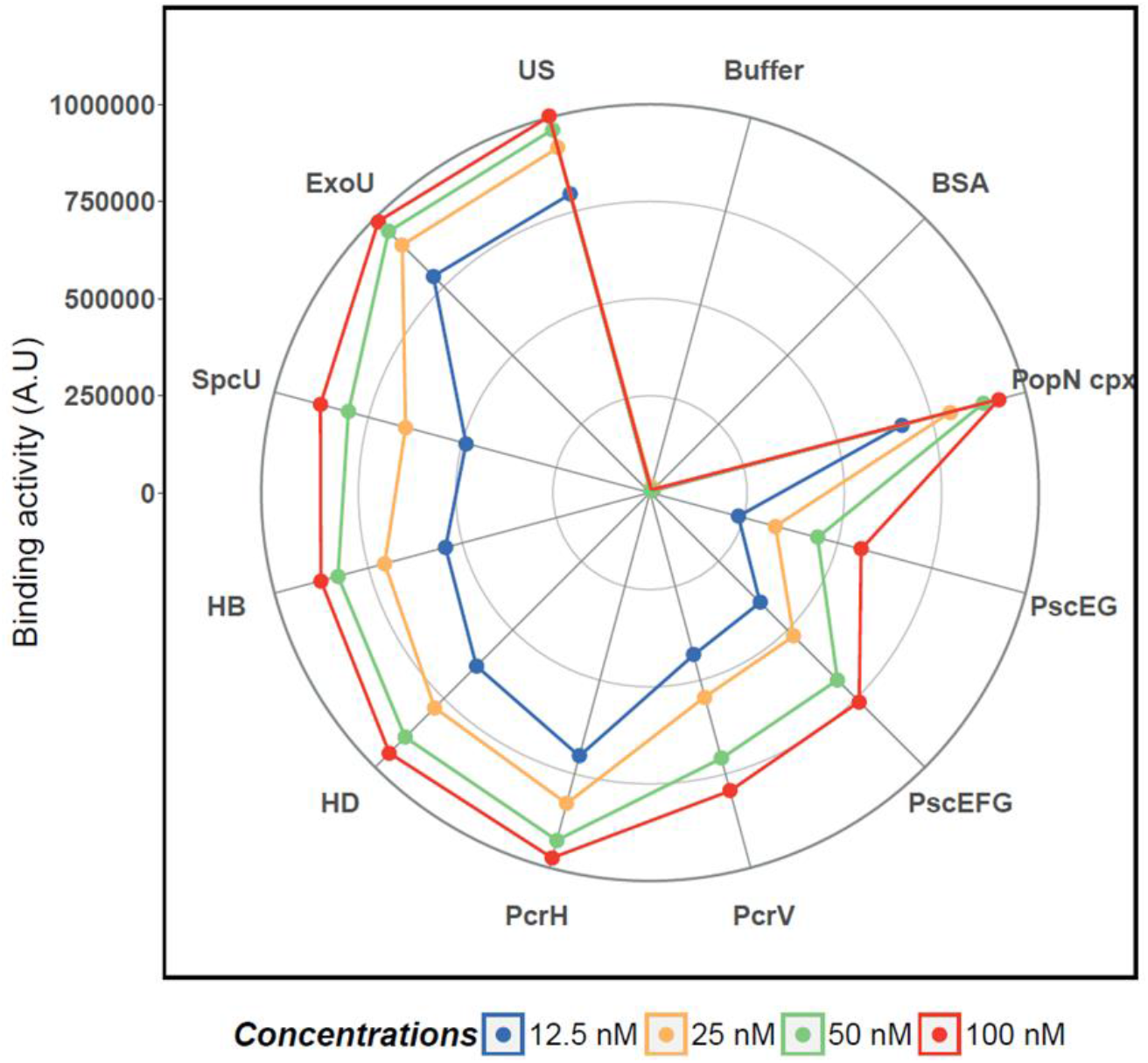
The ATPase PscN binds to T3SS soluble partners and their chaperones. The T3SS exported proteins, their chaperones alone or in complex were purified and 0.1 μM of each protein was coated and used in an ELISA assay to detect their interaction with the purified FSS-PscN at 12.5 nM, 25nM, 50 nM and 100 nM. BSA at 0.1 μM and PBS were used as negative controls. Individual proteins and complexes are: BSA (Bovine Serum Albumin), PopN cpx (PopN-Pcr1-Pcr2-PscB), PscEG (PscE-PscG), PscEFG (PscE-PscF-PscG), PcrV, PcrH, HD (PcrH-PopD), HB (PcrH-PopB), SpcU, ExoU, US (ExoU-SpcU).

To examine whether recombinant PscN could interact in solution with secreted proteins bound to their cognate chaperones, an HTRF assay was employed with four protein complexes including ExoU-SpcU (effector complex), PcrH-PopD (translocator complex), PscE-PscF-PscG (needle complex) and PopN-Pcr1-Pcr2-PscB (gate-keeper complex). HTRF relies on the detection of a FRET signal between a long-lasting lanthanide cryptate donor and an acceptor, allowing reduced interference from the buffer (Degorce et al., 2009). For this assay, we took advantage of the presence of a Flag-tag on PscN and a His-tag on its partners, enabling their labelling with antibodies directed toward these tags. Two pairs of donor and acceptor antibodies were used: anti-Flag-M2-Eu cryptate and anti-His-d2 or anti-Flag-M2-d2 and anti-His-Tb. However, only the interactions of PscN with the translocator and gate-keeper complexes could be detected (Figure S3). This lead to three hypotheses: i) the effector and needle complexes do not interact in solution with PscN; ii) the His-tags of effector and needle complexes are not accessible upon binding to PscN and iii) the binding site of the effector and needle complexes on PscN is different from that of the translocator and the gate-keeper complexes and the signal could consequently only be detected with those complexes that are close enough to the fluorophore-tagged N-terminus of PscN.

### PscN binds with a higher affinity to the effector complex

Another assay detecting protein binding in solution was required to investigate whether the absence of HTRF signal was artifactual. Therefore, microscale thermophoresis (MST) was employed to confirm the binding in solution and to determine the affinity between PscN and the exported proteins in *P. aeruginosa*. In these experiments, PscN was incubated with different concentrations of protein ligands and a modification of its thermophoresis signal indicated binding. A titration curve for each ligand, which represents the bound fraction depending on ligand concentration, was obtained and the determination of the dissociation constants (K_D_) was performed based on data from three independent replicate experiments. As shown by the binding curves, the interaction between PscN and the ExoU-SpcU effector complex displayed a high affinity (K_D_ = 45 ± 13 nM). In contrast, the measured affinities between PscN and the translocator complex PcrH-PopB/D (K_D_ = 4.7 ± 1.8 μM for PcrH-PopD and 4.9 ± 0.2 μM for PcrH-PopB) or the gate-keeper complex PopN-Pcr1-Pcr2-PscB (K_D_ = 0.8 ± 0.1 μM) were lower. The K_D_ between PscN and the needle complex (PscE-PscF-PscG) could not be determined due to a much lower affinity (predicted K_D_ > 150 μM) (Figure 3A and Table S2), suggesting that PscN probably did not interact with needle proteins in these conditions.

**Figure 3:**
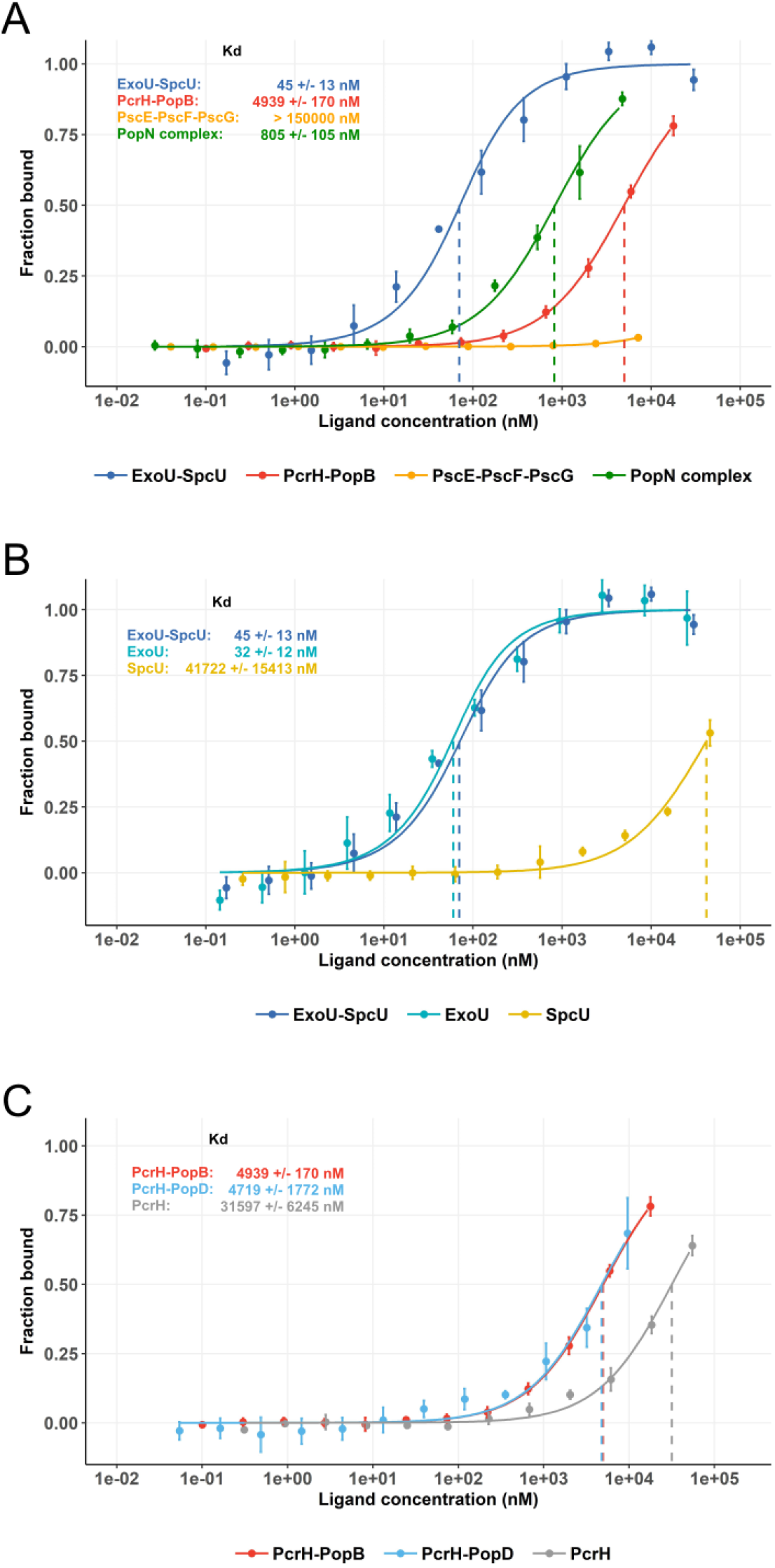
Microscale Thermophoresis allows the measurement of affinities between PscN and its T3SS soluble partners. FSS-PscN was labeled with MST dye and 100 nM of labeled protein were incubated with different concentrations of ligands. A titration fitted curve for each ligand, which represents the bound fraction depending on ligand concentrations, and K_D_ were obtained (A) Affinity between PscN and four complexes of effector, translocator, needle or gate-keeper. (B) Affinity between PscN and chaperone, effector or chaperone-effector complex. (C) Affinity between PscN and chaperone or chaperone-translocator complexes.

### PscN preferentially interacts with the protein complexes over chaperones alone

The SctN ATPase was shown to dissociate the cargo-chaperone complex by using energy from ATP hydrolysis in *S. typhimurium* (Akeda and Galan, 2005; Yoshida et al., 2014) and *Xanthomonas* (Lorenz and Buttner, 2009). It is likely that the cargo protein is subsequently secreted through the secretion channel while the chaperone is proposed to be released in the bacterial cytoplasm. To investigate this hypothesis, we examined the interaction preference of PscN for the complex, cargo or chaperone proteins. Through titration experiments using MST, the K_D_s between PscN and the effector ExoU or the cargo-chaperone complex ExoU-SpcU were measured to be 32 nM ± 12 nM and 45 nM ± 13 nM respectively. In contrast, the K_D_ between PscN and the chaperone alone was 1000 times higher, 42 ± 15 μM μM (Table S2 and Figure 3B). In similar experiments with the translocator chaperone PcrH bound or not to its cargos PopB and PopD, the K_D_s between the ATPase and the complex PcrH-PopB, 4.9 ± 0.2 μM, or PcrH-PopD, 4.7 ± 1.8 μM, were 6 fold-lower than the one between PscN and the chaperone PcrH alone (K_D_ = 32 ± 6 μM) (Figure 3C). Unfortunately, it is not possible to assess the K_D_ between PscN and PopB or PopD alone because these proteins are not stable *in vitro* without their cognate chaperone PcrH (Faudry et al., 2007; Wager et al., 2013). Nevertheless, taken together these results indicate that PscN would rather bind to cargo or complex proteins than to the chaperone protein alone. This supports the hypothesis that the chaperones are released in the bacterial cytoplasm after the complex dissociation.

### The gate keeper complex acts on PscN to regulate the binding of the needle complex protein to the ATPase

It has been shown that the gate-keeper protein SctW is involved in the secretion switch between translocator and effector proteins. A *ΔsctW* mutant strain was demonstrated to over-secrete the effectors in *P. aeruginosa, Shigella* and *Yersinia*. In contrast, the effect of this deletion mutation on translocator secretion is controversial (Ferracci et al., 2005; Roehrich et al., 2017; Yang et al., 2007). Interestingly, this protein SctW (MxiC), which is secreted by the T3SS, was shown to interact with the ATPase SctN (Spa47) in *Shigella* and the authors suggested that the gate-keeper inhibited the effector secretion by blocking its binding sites on the ATPase (Botteaux et al., 2009). As described in this work, ELISA, HTRF and MST assays showed that the PopN (SctW in *P. aeruginosa*) complex interacts with PscN. Therefore, to examine whether the PopN complex could play a role in the substrate sorting through binding to the ATPase, K_D_s were measured between the effector, translocator or needle complexes and PscN bound, or not bound, to the gate-keeper complex. For this purpose, PscN was previously incubated with the PopN complex at a concentration of 800 nM equal to the K_D_, yielding therefore half of the PscN population bound to the PopN complex. Unfortunately, it was not possible to use higher PopN complex concentration in this experiment because of the yield of its purification. Since only PscN is monitored by MST, a change in the apparent K_D_ would indicate an effect of the bound gate-keeper complex in the recognition capacities of the ATPase. While PscN alone almost does not bind to the needle complex (K_D_ estimated > 45 μM), the apparent K_D_ in presence of the gate-keeper was higher between PscN and PscE-PscF-PscG than between PscN and PcrH-PopB, 5.6 ± 2 μM and 12.4 ± 1 μM respectively (Table S2 and Figure 4). This result showed that binding of the gate-keeper to PscN dramatically modified its relative affinities for the translocator complex PcrH-PopB and the needle complex PscE-PscF-PscG, thus promoting the loading of the needle to PscN instead of the translocator. On the other hand, PopN complex had no effect on the binding of ExoU-SpcU to PscN (Figure 4), indicating that SctW does not block effector secretion through a direct competition for SctN binding, in contrast to the aforementioned hypothesis (Botteaux et al., 2009).

**Figure 4:**
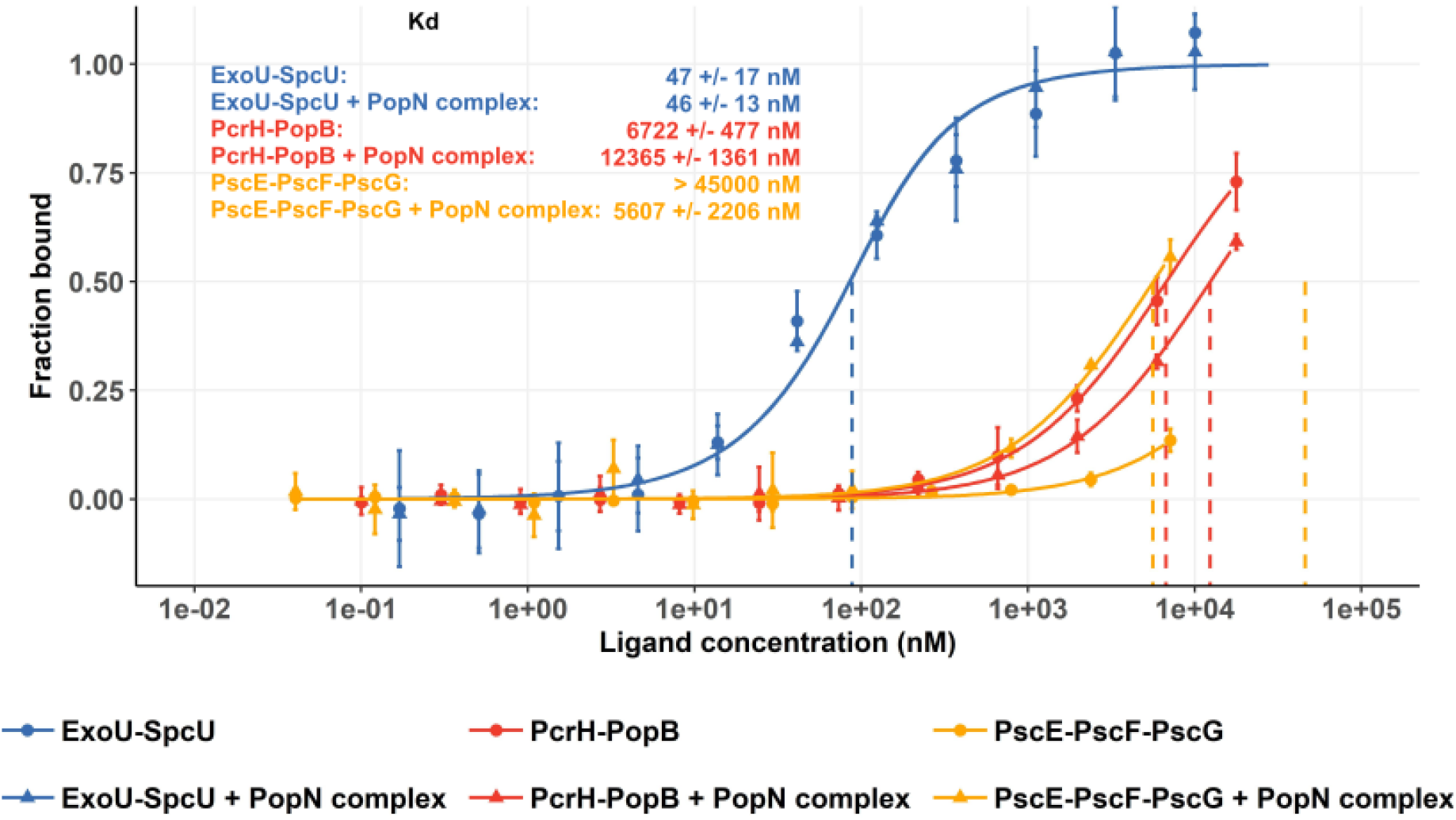
PopN complex modifies PscN affinity for the needle complex. FSS-PscN was labeled with MST dye and diluted at 100 nM in buffer containing or not PopN complex at 800nM. The mixtures were next incubated with different concentrations of three complexes: ExoU-SpcU, PcrH-PopB and PscE-PscF-PscG and a titration fitted curve of each condition was obtained. A significant shift of apparent K_D_ between PscN and PscE-PscF-PscG in presence of PopN complex indicates that the gate-keeper plays a role in the loading of needle complex on the ATPase.

However, it would be possible that a direct interaction between the PscF complex and the PopN complex bound to PscN would be responsible for the apparent increase of the affinity between PscN the needle complex protein. To rule out this possibility, SEC analysis of the individual complexes and a mixture of the PscF and PopN complexes were performed. The superposition of the chromatograms clearly shows that there was no interaction between the needle and gate-keeper complexes while this assay was performed at protein concentration close to the highest concentration used in MST (Figure S4). Furthermore, the PscF complex was labeled with MST dye and thermophoresis experiments were conducted with increasing concentrations of PopN complex. No interaction could be detected with PopN complex concentration up to 15 μM, thus confirming the absence of interaction between these two complexes. Overall, this work suggests a new role of the gate-keeper and the ATPase in the control of substrate hierarchical secretion in *P. aeruginosa*.

## Discussion

The ATPase SctN is essential for the assembly and function of the T3SS. It was first described to interact with exported proteins in *Salmonella, Escherichia coli, Shigella* and *Xanthomonas* species (Akeda and Galan, 2005; Akeda and Galán, 2004; Allison et al., 2014; Botteaux et al., 2009; Cooper et al., 2010; Gauthier, A. and Finlay, 2003; Lorenz and Buttner, 2009; Yoshida et al., 2014; Zarivach et al., 2007). However, an extensive comparison of the interactions between the ATPase and the secreted proteins remained to be performed to get insight into its function.

To date, only structures of the N-terminally truncated monomers of SctN were obtained by X-ray crystallography (Allison et al., 2014; Bernal et al., 2019; Burgess et al., 2016; Zarivach et al., 2007), suggesting that the N-terminal domain of SctN is flexible and/or partially unfolded. Some groups were able to heterologously express and purify the His-tagged recombinant full-length protein from *E. coli* (Akeda and Galan, 2005; Andrade et al., 2007; Chatterjee et al., 2013; Yoshida et al., 2014). Unfortunately, we were not able to obtain PscN in the same production condition despite many attempts. This indicates that although SctN shares 40-50% of sequence identity among the Gram-negative pathogenic bacteria (Zarivach et al., 2007), the T3SS ATPase of *P. aeruginosa* is more difficult to produce in *E. coli*. Indeed, a recent study of PscN required refolding after solubilisation of inclusion bodies (Halder et al., 2018). Therefore, in order to obtain the purified protein, we fused PscN to a triple tag and achieved efficient production and purification under native conditions from an engineered *P. aeruginosa* strain (Arora et al., 1997; Brunschwig and Darzins, 1992). Consequently, we could obtain recombinant full-length PscN in sufficient quantities for biochemical analysis. Furthermore, this purified PscN is catalytically active and the relatively long tag at the N-terminus is not deleterious for its activity *in vivo*.

By using a protein-protein interaction ELISA assay, we demonstrated that PscN interacts with the chaperones, cargos and complexes of effector, translocator, needle and gate-keeper proteins. These results were next assessed by MST and the affinity between PscN and the four complexes was determined. Interestingly, PscN preferentially interacts with chaperone-cargo complexes compared to chaperone proteins alone. Thus, this lower affinity for the chaperones could promote their release in the bacterial cytoplasm after the dissociation from their cargo proteins upon ATP hydrolysis. Actually, the binding of ATP to SctN induces a conformation change of the C-terminal domain which was proposed to be the binding site for effector-chaperone complex on SctN (Gao et al., 2018; Majewski et al., 2019). However, the presence of ATP did not affect the affinity of PscN for the effector complex ExoU-SpcU in *P. aeruginosa* (Figure S5).

Using structural modeling *in silico*, chaperones of effectors were docked to the loop of a two-helix-finger motif in the C-terminal of SctN - SsaN in *S. enterica* and EscN in enteropathogenic *E. coli* (Allison et al., 2014; Zarivach et al., 2007) and the mutation of a conserved residue (V379P) in this region abolished the interaction of the ATPase SsaN and the effector chaperone SrcA in *S. enterica* SPI-2 *in vitro* (Allison et al., 2014). Interestingly, the corresponding mutation (L376P) was also identified from the screening of loss-of-function mutations of the ATPase InvC in *S. enterica* SPI-1 and affected the secretion of the early substrate InvJ (SctP), middle substrates SipB/C (SctE/B) and the effector SptP (Akeda and Galán, 2004). Hence, this two-helix-finger region was pointed out as the binding site of SctN to exported proteins in T3SS and these proteins could compete to dock to this site on the ATPase. In this case, it would be tempting to speculate that different affinities would dictate the order of protein secretion. Nonetheless, the affinity between PscN and translocator (K_D_ in the 5 μM range) and effector (K_D_ in the 50 nM range) seems contradictory to the hierarchical secretion of these substrates because the translocators are secreted prior to the effectors. Actually, an interpretation solely based on differences in affinities would be too simplistic. First, because the local concentrations of the different PscN partners in bacteria are extremely challenging to determine and second because these partners are continually expressed after T3SS activation and a significant decrease of their concentrations during T3SS secretion is speculative. On the other hand, PscN bound to either ExoU-SpcU (Effector-Chaperone) or PcrH-PopB (Translocator-Chaperone) had almost the same affinity for each substrate as the free-PscN (Figure S6). Therefore, late and middle substrates seem not to compete with each other to dock to a same site on PscN.

The existence of different binding sites of substrates on SctN could be supported by the fact that no inhibition of binding between PscN and ExoU-SpcU, PcrH-PopB or PscE-PscF-PscG (late, middle and early substrates) when PscN was bound to the gate-keeper (SctW) PopN complex. In contrast, binding of PopN complex to PscN promoted an increase of the affinity between PscN and PscE-PscF-PscG. We therefore propose a model of allosteric conformational change of PscN that regulates its recognition of the needle protein complex (Figure 5). This allosteric regulation of SctN would participate in the mechanism of the T3SS secretion switch from early to middle substrate, which also involves SctP and SctU but that are still generally unclear.

**Figure 5:**
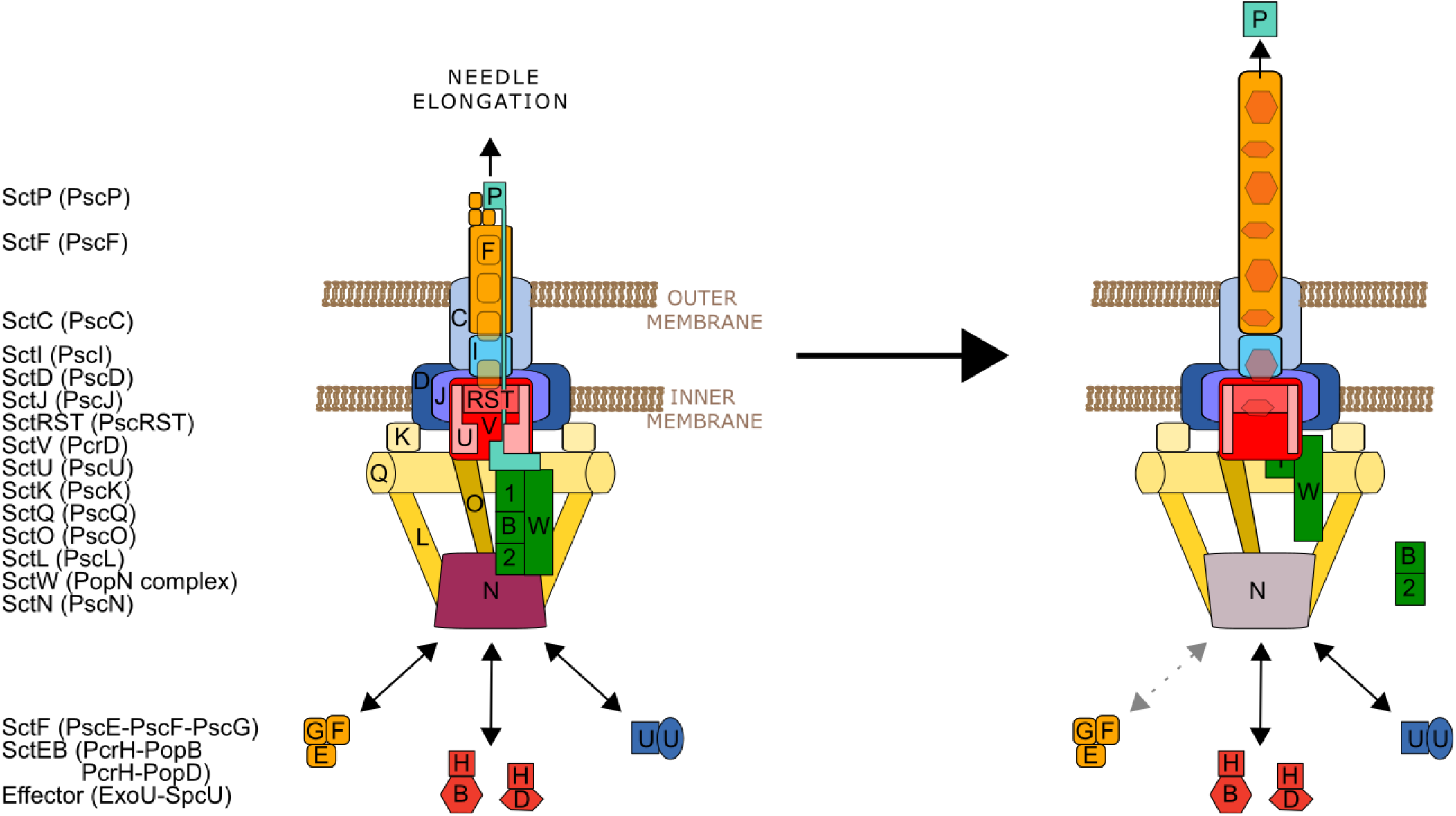
Model of allosteric conformation change of the ATPase PscN controlling the needle protein export. (1) In the presence of the SctW complex, SctN is in the “needle binding on” state which facilitates its interaction with the needle complex and SctU prevents the loading of the translocators SctE and SctB and effectors. The needle subunit SctF is therefore secreted to form the T3SS needle while its chaperones (PscE and PscG in *P. aeruginosa*) are released in the cytoplasm. (2) When the needle reaches its final length, controlled by SctP, SctP is secreted, SctU undergoes auto-proteolysis and SctW is detached from SctN and tethered to SctV. The dissociation of SctW from SctN induce a SctN conformational change preventing its interaction with the needle complex. The needle subunit could not be recruited to the export gate anymore. PscU auto-cleavage allows the secretion of translocators while their chaperone (PcrH in *P. aeruginosa*) is released in the cytoplasm. A further switch not depicted here prevents translocator secretion and allows effector injection into the host cell. In *P. aeruginosa* the proteins are: SctW complex PopN-Pcr1(1)-Pcr2(2)-PscB(B); SctF needle complex: PscE-PscF-PscG; SctE and SctB translocators in complexes with their chaperone: PcrH-PopB/D; Effector complex: ExoU-SpcU.

The length of the T3SS injectisome needle is controlled by the ruler SctP that is strongly conserved with a “ball and chain” architecture. This protein family consists of a C-terminal compactly folded domain (the ball) followed by a C-terminus disordered tail, an N-terminal disordered ruler domain (the chain) and a flexible linker connecting these two domains (Bergeron et al., 2016; Kinoshita et al., 2020; Minamino, 2018). Another regulator, the autoprotease SctU, consists of an N-terminal transmembrane domain and a C-terminal cytoplasmic domain, which undergoes auto-cleavage at the conserved NPTH loop, allowing the secretion of translocators and effectors (Frost et al., 2012; Sorg et al., 2007; Wood et al., 2008). The C-terminal of SctU was shown to bind to SctP at both N-terminal domain (at an export signal in the firsts 40 amino acids) and to the flexible linker (in T3SS) or to the C-terminal folded domain (in the related flagellum system) (Bergeron et al., 2016; Ho et al., 2017; Kinoshita et al., 2017, 2020; Minamino et al., 1999). Despite this interaction, the arrest of needle elongation and the switching to middle substrate secretion might be two separate events because some mutations preventing auto-cleavage of SctU abolished secretion of middle and late substrates but did not affect the control of early substrate secretion and the needle length (Monjarás Feria et al., 2015; Wagner et al., 2018; Wood et al., 2008). One element linking the needle length measurement to the termination of the secretion of the needle subunit is therefore lacking. Indeed, focused on exported substrates recognition by the T3SS injectisome before secretion, our findings give new insights on the regulation of needle assembly.

We propose a new function of the gate-keeper SctW complex, which would act on the interaction of SctN with the SctF needle complex. In the resting state before the beginning of the T3SS secretion process, we propose that the SctW complex is recruited to the T3SS sorting platform through binding to SctN and to SctU-SctP at the same time (Figure 5A). Indeed, the ruler SctP was shown to directly interact with the gate-keeper SctW (Shaulov et al., 2017). This SctW-SctN interaction would allow needle complex recognition by enabling its interaction with the ATPase while SctU would prevent translocators and effectors to be loaded into the T3SS. When the T3SS needle reaches the expected length, fully stretched SctP would promote its dissociation from SctW and SctU and its secretion, which in turn modulates the rate of SctU auto-proteolysis (Ho et al., 2017) facilitating the secretion of translocators and effectors. As already shown, the gate-keeper SctW can also be tethered to the inner-membrane protein SctV (Gaytán et al., 2018; Lee et al., 2014; Yu et al., 2018). There, it can fulfill its second function: the regulation of the next secretion switch, from middle to late substrates, which may also involve the sorting platform (Lara-Tejero et al., 2011). We thus suppose that following the secretion of SctP and the auto-proteolysis of SctU, the SctW complex would be displaced from the SctN ATPase to the export gate protein SctV (Figure 5B). While it is plausible that PscB and Pcr2 chaperones could be dissociated from the SctW complex by SctN activity, chaperone dissociation from secreted proteins by SctN has so far only been demonstrated for translocators and effectors (Akeda and Galan, 2005; Lorenz and Buttner, 2009; Yoshida et al., 2014). Freed from SctW, SctN affinity for the needle complex would drastically drop down and this complex would not be handled anymore, preventing further secretion of needle subunits.

Localized at the cytoplasmic extremity of the apparatus, the T3SS ATPase may be the first platform that screens and loads exported substrates to the T3SS. The loaded proteins would be next secreted in a timing that integrates signals from regulators such as the ruler SctP, the switch SctU and/or the gate-keeper SctW. Of importance, our findings reveal a new role of the gate-keeper complex in the control of needle subunit secretion, thus complementing an overview of the mechanisms by which the secretion switches from early to middle substrates.

## Materials and methods

### Bacterial strains and plasmids

The *E. coli* and *P. aeruginosa* strains and plasmids used in this study are listed in the Table 4. The *P. aeruginosa* deletion strain *ΔpscN* was constructed using the method of Slice Overlapping Extension PCR (SOE-PCR). The Strep-tagged PscN construction was generated by PCR using primers NdeI-Strep-PscN and PscN-HindIII (Table S1) and the amplified fragments were cloned into the pIApG plasmid. To generate the Flag-Strep-Strep-PscN construct, a DNA fragment containing NcoI-Flag-Strep-Strep-NterminusPscN-NotI synthesized by Invitrogen, was cloned into the NcoI/NotI sites of pIApG-Strep-PscN. Both constructions pIApG-Strep-PscN and pIApG-FSS-PscN were transformed into *P. aeruginosa* CHA *ΔpscN*. Otherwise, the genes encoding Strep-PscN and FSS-PscN were cloned from pIApG to pET15b or pET15bVP vectors using NcoI/BamHI sites. The pET15b-Strep-PscN and pET15b-FSS-PscN were next transformed into *E. coli* BL21 (DE3) while the pET15bVP-FSS-PscN was transformed into *P. aeruginosa* PAO1 ADD1976 (Arora et al., 1997; Brunschwig and Darzins, 1992). To generate the PopN complex expression construct, popN pcr1 pcr2 were amplified from the chromosome by PCR, using primers PopNEX5 and Pcr2fB-3. This PCR product was then fused to pscB by amplifying pscB using primers PscBf2-5 and PscB3H and combining the resultant PCR product to the popNpcr1pcr2 PCR product using splicing by overlap extension PCR. This product was then cloned into pET28bTEV (Classen et al., 2003) as a NdeI/HindIII restriction fragment to give pET28bTEV-His-PopN-Pcr1-Pcr2-PscB

### LDH release assay

Macrophages from the J774 cell line were grown overnight in a 96 well plate at a density of 50,000 cells per well. *P. aeruginosa* CHA WT, *ΔpscN*, and *ΔpscN*/pIApG-FSS-PscN were grown at 37°C in LB medium to OD 1 and added to the J774 cells at a multiplicity of infection (MOI) of 10. Cell death was monitored at 2 h, 3 h and 4 h post-infection using a cytotoxicity detection kit (lactate dehydrogenase (LDH); Roche). The assay was performed in triplicate.

### Protein expression and purification

For pull-down experiments of PscN-partners, *P. aeruginosa* CHA harboring pIApG-Strep-PscN was grown at 37°C in LB medium containing 20 mM of MgCl2 and 5 mM of EGTA to OD 1. Harvested cells were resuspended in buffer containing 100 mM Tris pH 8, 150 mM NaCl and 1 mM EDTA. Cells were lysed by sonication and 1% triton X100 was added to solubilize membrane fraction for 14 h at 4°C. The non-soluble membranes were eliminated by ultracentrifugation at 125,000 g for 1h30 at 4°C and PscN was purified from the supernatant using a Streptrap HP column on an Akta purifier system. The purified Strep-PscN fractions were eluted in buffer containing 2.5 mM desthiobiotin and next analyzed by mass spectrometry.

*E. coli* BL21(DE3) harboring Strep-PscN and FSS-PscN and PAO1 ADD1976 harboring FSS-PscN were grown at 37°C in 500 ml of LB medium with appropriate antibiotic and were induced with 1 mM IPTG at OD 1 for 2 h at 28°C. Cells were harvested and resuspended in 20 ml of binding buffer containing 50 mM Tris pH 9, 50 mM Arginine, 50 mM Glutamate, 150 mM NaCl, 10% Glycerol, 5 mM TCEP and 1% Triton X100. Cell suspension was lysed by sonication and centrifuged at 200,000 g for 30 minutes. Supernatants were next incubated with Streptactin sepharose beads for 15 h at 4°C and consecutively washed with binding buffer containing 1% and 0.1% of triton X100. Proteins were then eluted with 2.5 mM desthiobiotin.

*E. coli* BL21 (DE3) harboring expression vectors for 6His-tagged ExoU-SpcU, ExoU, SpcU, PcrH-PopB, PcrH-PopD, PcrH, PcrV, PscEFG, PscE-PscG, PopN complex were grown at 37°C in 500 ml of LB medium with appropriate antibiotic and were induced with 1 mM IPTG at OD 0.6 for 3 h. Harvested cells were resuspended in 20 ml of IMAC buffer (25 mM Tris pH 8, 500 mM NaCl) containing 10 mM Imidazole. Cell suspension was lysed by sonication and centrifuged at 200,000 g for 30 minutes. Soluble proteins were purified using Histrap HP column on an Akta purifier system. The proteins were eluted by the IMAC buffer supplemented with 200 mM Imindazole and subsequently injected onto a Supedex 200 Increase size exclusion column pre-equilibrated with a buffer containing 50 mM Tris pH 8 and 150 mM NaCl. All the proteins displayed the expected elution profiles.

### Protein quantification and ATPase activity

The fractions of all the purified protein were quantified on Agilent Bio-analyzer chips. Samples were prepared according to the instruction of the Agilent protein 80 kit and analyzed on a 2100 Bio-analyzer device (Agilent technologies). The ATPase activity of FSS-PscN was determined by measuring the release of total phosphate at 37°C for 30 minutes which was detected by the malachite green method (Casabona et al., 2013a).

### MS-based proteomic analyses

Proteins extracted from pull-down eluates were stacked in the top of a SDS-PAGE gel (NuPAGE 4-12%, ThermoFisher Scientific), stained with Coomassie blue (R250, Bio-Rad) before being in-gel digested using modified trypsin (Promega, sequencing grade) as previously described (Casabona et al., 2013b). Of note, samples from Streptactin and His-tag pull-downs were prepared using respectively oxidation and reduction-alkylation procedures (Jaquinod et al., 2012). Resulting peptides were analyzed by online nanoLC-MS/MS (respectively UltiMate 3000 and LTQ-Orbitrap XL and UltiMate 3000 RSLCnano and Q-Exactive HF, Thermo Scientific, for Streptactin and His-tag pull-down eluates). For this, peptides were sampled on a 300 μm x 5 mm PepMap C18 precolumn (Thermo Scientific) and separated on 75 μm x 150 mm and 75 μm x 250 mm C18 columns (Gemini C18, 3 μm, Phenomenex, and Reprosil-Pur 120 C18-AQ, 1.9 μm, Dr. Maisch). MS and MS/MS data were acquired using Xcalibur (Thermo Scientific). Mascot Distiller (Matrix Science) was used to produce mgf files before identification of peptides and proteins using Mascot (version 2.7) through concomitant searches against Uniprot (*P. aeruginosa* PAO1 taxonomy for Streptactin pull-down eluate and *E. coli* BL21 strain for His-tag pull-down eluate, April 2020 versions), specific sequences of tagged and co-expressed proteins (homemade), classical contaminants database (homemade) and the corresponding reversed databases. The Proline software (Bouyssié et al., 2020) was used to filter the results (conservation of rank 1 peptides, peptide length ≥ 6 amino acids, peptide-spectrum match identification FDR < 1% as calculated on peptide scores by employing the reverse database strategy, minimum peptide-spectrum-match score of 25, and minimum of 1 specific peptide per identified protein group) and extract the quantification information.

### ELISA

50 μl of each tested proteins ExoU-SpcU, ExoU, SpcU, PcrH-PopB, PcrH-PopD, PcrH, PcrV, PscEFG, PscE-PscG, PopN complex diluted in PBS at 0.1 μM, were coated on ELISA 96 well plates for 15 h at 4°C. The BSA diluted in PBS at 0.1 μM and PBS buffer were used as negative controls. The wells were then washed three times with 200 μl of PBS/0.1% Tween 20 (PBST) and incubated with 200 μl of PBS containing BSA 4% for 4 h. The solution was next removed and plates were washed three times. Afterward, 50 μl of purified FSS-PscN diluted in PBST containing BSA 4% (PBST.BSA 4%) at 12. nM, 25 nM, 50 nM and 100 nM, were incubated for 1 h at room temperature (RT°). The wells were subsequently washed three times with 200 μl of PBST and 50 μl of anti-PscN antibody (1/2000 in PBST.BSA 4%) were incubated for 1 h at RT°. Three additional washes with 200 μl of PBST were performed before 50 μl of anti-rabbit-HRP antibody (1/40000 in PBST.BSA 4%) were added for 1 h at RT°. The wells were then washed with PBST (three time) and revealed using the Pierce ECL Western Blotting Substrate (ThermoFisher: 32106). The chemi-luminescence was measured using a FluoStar BMG reader.

### Two-hybrid analysis

To generate cI fusions to *popN* or *pscB*, the open reading frames were amplified (*popN*: PopN-5NotI + PopN-3AscI, *pscB*: PscB-5NotI + PscB-3AscI,) and cloned as NotI-AscI fragments into plasmid pACλcI35 (Dove and Hochschild, 2004). Fusions to RNA polymerase α were generated by amplifying *pcr1, pcr2*, or *pscB* (*pscB*: as for cI, *pcr1*: Pcr1-5NotI + Pcr1-3AscI, *pcr2*: Pcr2-5NotI + Pcr2-3AscI) and cloning the resultant PCR products as NotI-AscI fragments into plasmid pBRα35 (Dove and Hochschild, 2004). To co-express *pscB* with the α-pcr2 fusion gene, *pscB* was cloned in a translationally coupled manner downstream of *pcr2* in pBRa-pcr2 (primers: PscBf2-5 + Pcr2fB-3). To assay interactions, plasmids expressing fusion genes to cI and α, respectively, were co-transformed into strain BN469 (Nickels, 2009), which harbors the test promoter that includes a cI operator site, and in which lacZ expression is activated by recruiting RNA polymerase via an interaction between the cI and α fusion proteins. Three individual colonies were assayed for each plasmid combination. Where indicated, production of the fusion proteins was induced with 50 μM IPTG. Activation of the test promoter was assayed by β-galactosidase assay, as previously described (Miller, 1992).

### HTRF

Purified FSS-PscN, ExoU-SpcU, PcrH-PopD, PscE-PscF-PscG and PopN complexes were diluted in PBS/0.1% Tween 20 containing BSA 0.05% (PBST.BSA 0.05%) at the indicated concentrations. The fluorophore-conjugated antibodies Flag-d2, His-Tb, His-d2 were diluted in PBST.BSA 0.05% while Flag-Eu^3+^ was diluted in PBST.BSA 0.05% containing 400 mM KF. The concentration of fluorophore-conjugated antibodies was adjusted following the recommendation from the manufacturer (Cisbio). Experiments were performed in 384 well plates with a final volume of 20 μl per well. Reagents were sequentially added: 5 μl PscN, 5 μl ligand protein and 5 μl anti-Flag-Eu^3+^/anti-His-d2 or anti-His-Tb/anti-Flag-d2. After 6 h of incubation at room temperature, the fluorescence was measured on an Infinite M1000 Tecan reader.

### Microscale thermophoresis (MST)

Purified FSS-PscN fractions were buffer-exchanged to a buffer containing 50 μM HEPES pH 8, 150 mM NaCl, 10% glycerol and 0.1% triton X100 by using a desalting column PD Spin Trap G25 (GE healthcare: 28-9180-04). This FSS-PscN solution was then labeled with the Monolith Protein Labeling Kit RED-NHS (MO-L001) following the manufacturer instruction (Nanotemper) and diluted in the purification binding buffer (see above) containing 0.1% triton X100 and without TCEP. For MST measurements, labeled FSS-PscN was diluted at 100 nM in the buffer containing 50 mM Tris pH 6.7, 50 mM NaCl, 5 mM DTT and 0.1% Pluronic F127. The ligand proteins were prepared at the indicated concentrations in SEC buffer containing 50 mM Tris pH 8 and 150 mM NaCl. PscN and the ligand were mixed in 1:1 v/v ratio, resulting in a final buffer which contains 50 mM Tris pH 7.1, 100 mM NaCl, 2.5 mM DTT and 0.05% Pluronic F127. The thermophoresis signal was then measured on a Monolith NT.115 device (Nanotemper) with premium capillaries (MO-K025). Data from three independent replicates were used to estimate the K_D_ using the MO.Affinity Analysis software provided by the manufacturer Nanotemper.

For three-partners assay, the same experiments were performed except that FSS-PscN diluted in 50 mM Tris pH 6.7, 50 mM NaCl, 5 mM DTT and 0.1% Pluronic F127 was mixed with first partner diluted in 50 mM Tris pH 8, 50 mM NaCl, 5 mM DTT and 0.1% Pluronic F127. This protein mix containing PscN at 100 μM and the first partner was then added (1:1 v/v) to the ligand proteins diluted in 50 mM Tris pH 8 and 150 mM NaCl. The final buffer composition was therefore different from the one described above for the two-partners assay.

## Acknowledgements

This work was supported by the associations “Vaincre la mucoviscidose” and “Gregory Lemarchal”, grants from the AVIESAN T3SS (ANR PRP1.4), the Laboratory of Excellence GRAL (ANR-10-LABX-49-01) and the Agence Nationale de Recherche (ANR-15-CE11-0018-01). Proteomic experiments were partly supported by the Agence Nationale de la Recherche (ProFI grant ANR-10-INBS-08-01). Authors acknowledge the platforms supported by GRAL, financed within the University Grenoble Alpes graduate school (Ecoles Universitaires de Recherche) CBH-EUR-GS (ANR-17-EURE-0003). We are grateful to Caroline Mas and Cécile Morlot for help with the Microscale Thermophoresis experiments and we sincerely acknowledge the help, support and critical reading of Andréa Dessen.

## CRediT author statement

**Tuan-Dung Ngo:** Conceptualization, Formal analysis, Investigation, Writing - Original Draft, Visualization. **Caroline Perdu:** Conceptualization, Investigation. **Bakhos Jneid:** Methodology, Investigation. **Michel Ragno:** Investigation. **Julia Novion Ducassou:** Investigation. **Alexandra Kraut:** Investigation. **Yohann Couté:** Formal analysis, Writing - Review & Editing, Visualization, Supervision. **Charles Stopford:** Investigation. **Ina Attree:** Conceptualization, Writing - Review & Editing, Supervision, Funding acquisition. **Arne Rietsch:** Conceptualization, Writing - Review & Editing, Visualization, Supervision. **Eric Faudry:** Conceptualization, Investigation, Writing - Original Draft, Visualization, Supervision, Project administration, Funding acquisition.

## References

Akeda, Y., and Galán, J.E. (2004). Genetic analysis of the *Salmonella enterica* type III secretion-associated ATPase InvC defines discrete functional domains. J. Bacteriol. 186, 2402–2412.

Akeda, Y., and Galan, J.E. (2005). Chaperone release and unfolding of substrates in type III secretion. Nature 437, 911–915.

Allison, S.E., Tuinema, B.R., Everson, E.S., Sugiman-Marangos, S., Zhang, K., Junop, M.S., and Coombes, B.K. (2014). Identification of the Docking Site between a Type III Secretion System ATPase and a Chaperone for Effector Cargo. J. Biol. Chem. 289, 23734–23744.

Andrade, A., Pardo, J.P., Espinosa, N., Pérez-Hernández, G., and González-Pedrajo, B. (2007). Enzymatic characterization of the enteropathogenic *Escherichia coli* type III secretion ATPase EscN. Arch. Biochem. Biophys. 468, 121–127.

Arora, S.K., Ritchings, B.W., Almira, E.C., Lory, S., and Ramphal, R. (1997). A transcriptional activator, FleQ, regulates mucin adhesion and flagellar gene expression in *Pseudomonas aeruginosa* in a cascade manner. J. Bacteriol. 179, 5574–5581.

Belyy, A., Mechold, U., Renault, L., and Ladant, D. (2018). ExoY, an actin-activated nucleotidyl cyclase toxin from <i>P. aeruginosa<i>: A minireview. Toxicon Off. J. Int. Soc. Toxinology 149, 65–71.

Bergeron, J.R.C., Fernández, L., Wasney, G.A., Vuckovic, M., Reffuveille, F., Hancock, R.E.W., and Strynadka, N.C.J. (2016). The Structure of a Type 3 Secretion System (T3SS) Ruler Protein Suggests a Molecular Mechanism for Needle Length Sensing. J. Biol. Chem. 291, 1676–1691.

Bernal, I., Römermann, J., Flacht, L., Lunelli, M., Uetrecht, C., and Kolbe, M. (2019). Structural analysis of ligand-bound states of the *Salmonella* type III secretion system ATPase InvC. Protein Sci. Publ. Protein Soc. 28, 1888–1901.

Botteaux, A., Sory, M.P., Biskri, L., Parsot, C., and Allaoui, A. (2009). MxiC is secreted by and controls the substrate specificity of the *Shigella flexneri* type III secretion apparatus. Mol. Microbiol. 71, 449–460.

Bouyssié, D., Hesse, A.-M., Mouton-Barbosa, E., Rompais, M., Macron, C., Carapito, C., Gonzalez de Peredo, A., Couté, Y., Dupierris, V., Burel, A., et al. (2020). Proline: an efficient and user-friendly software suite for large-scale proteomics. Bioinformatics 36, 3148–3155.

Brunschwig, E., and Darzins, A. (1992). A two-component T7 system for the overexpression of genes in *Pseudomonas aeruginosa*. Gene 111, 35–41.

Burgess, J.L., Burgess, R.A., Morales, Y., Bouvang, J.M., Johnson, S.J., and Dickenson, N.E. (2016). Structural and Biochemical Characterization of Spa47 Provides Mechanistic Insight into Type III Secretion System ATPase Activation and *Shigella* Virulence Regulation. J. Biol. Chem. jbc.M116.755256.

Casabona, M.G., Silverman, J.M., Sall, K.M., Boyer, F., Couté, Y., Poirel, J., Grunwald, D., Mougous, J.D., Elsen, S., and Attree, I. (2013a). An ABC transporter and an outer membrane lipoprotein participate in posttranslational activation of type VI secretion in *Pseudomonas aeruginosa*. Environ. Microbiol. 15, 471–486.

Casabona, M.G., Vandenbrouck, Y., Attree, I., and Couté, Y. (2013b). Proteomic characterization of *Pseudomonas aeruginosa* PAO1 inner membrane. PROTEOMICS 13, 2419–2423.

Chatterjee, R., Halder, P.K., and Datta, S. (2013). Identification and Molecular Characterization of YsaL (Ye3555): A Novel Negative Regulator of YsaN ATPase in Type Three Secretion System of Enteropathogenic Bacteria *Yersinia enterocolitica*. PLoS ONE 8, e75028.

Chen, L., Ai, X., Portaliou, A.G., Minetti, C.A.S.A., Remeta, D.P., Economou, A., and Kalodimos, C.G. (2013). Substrate-Activated Conformational Switch on Chaperones Encodes a Targeting Signal in Type III Secretion. Cell Rep. 3, 709–715.

Classen, S., Olland, S., and Berger, J.M. (2003). Structure of the topoisomerase II ATPase region and its mechanism of inhibition by the chemotherapeutic agent ICRF-187. Proc. Natl. Acad. Sci. 100, 10629–10634.

Cooper, C.A., Zhang, K., Andres, S.N., Fang, Y., Kaniuk, N.A., Hannemann, M., Brumell, J.H., Foster, L.J., Junop, M.S., and Coombes, B.K. (2010). Structural and Biochemical Characterization of SrcA, a Multi-Cargo Type III Secretion Chaperone in *Salmonella* Required for Pathogenic Association with a Host. PLoS Pathog. 6, e1000751.

Degorce, F., Card, A., Soh, S., Trinquet, E., Knapik, G.P., and Xie, B. (2009). HTRF: a technology tailored for drug discovery–a review of theoretical aspects and recent applications. Curr. Chem. Genomics 3, 22.

Demler, H.J., Case, H.B., Morales, Y., Bernard, A.R., Johnson, S.J., and Dickenson, N.E. (2019). Interfacial amino acids support Spa47 oligomerization and *Shigella* Type Three Secretion System activation. Proteins 87, 931–942.

Deng, W., Marshall, N.C., Rowland, J.L., McCoy, J.M., Worrall, L.J., Santos, A.S., Strynadka, N.C.J., and Finlay, B.B. (2017). Assembly, structure, function and regulation of type III secretion systems. Nat. Rev. Microbiol. 15, 323–337.

Diepold, A., and Wagner, S. (2014). Assembly of the bacterial type III secretion machinery. FEMS Microbiol. Rev. 38, 802–822.

Discola, K.F., Förster, A., Boulay, F., Simorre, J.-P., Attree, I., Dessen, A., and Job, V. (2014). Membrane and chaperone recognition by the major translocator protein PopB of the type III secretion system of *Pseudomonas aeruginosa*. J. Biol. Chem. 289, 3591–3601.

Dove, S.L., and Hochschild, A. (2004). A Bacterial Two-Hybrid System Based on Transcription Activation. In Protein-Protein Interactions, (New Jersey: Humana Press), pp. 231–246.

Edqvist, P.J., Olsson, J., Lavander, M., Sundberg, L., Forsberg, A., Wolf-Watz, H., and Lloyd, S.A. (2003). YscP and YscU regulate substrate specificity of the *Yersinia* type III secretion system. J. Bacteriol. 185, 2259–2266.

Erhardt, M., Mertens, M.E., Fabiani, F.D., and Hughes, K.T. (2014). ATPase-independent type-III protein secretion in Salmonella enterica. PLoS Genet. 10, e1004800.

Faudry, E., Job, V., Dessen, A., Attree, I., and Forge, V. (2007). Type III secretion system translocator has a molten globule conformation both in its free and chaperone-bound forms. FEBS J. 274, 3601–3610.

Ferracci, F., Schubot, F.D., Waugh, D.S., and Plano, G.V. (2005). Selection and characterization of *Yersinia pestis* YopN mutants that constitutively block Yop secretion. Mol. Microbiol. 57, 970–987.

Frost, S., Ho, O., Login, F.H., Weise, C.F., Wolf-Watz, H., and Wolf-Watz, M. (2012). Autoproteolysis and intramolecular dissociation of *Yersinia* YscU precedes secretion of its C-terminal polypeptide YscU(CC). PloS One 7, e49349.

Gao, X., Mu, Z., Yu, X., Qin, B., Wojdyla, J., Wang, M., and Cui, S. (2018). Structural Insight Into Conformational Changes Induced by ATP Binding in a Type III Secretion-Associated ATPase From *Shigella flexneri.* Front. Microbiol. 9.

Gauthier, A., and Finlay, B.B. (2003). Translocated Intimin Receptor and Its Chaperone Interact with ATPase of the Type III Secretion Apparatus of Enteropathogenic *Escherichia coli.* J. Bacteriol. 185, 6747–6755.

Gaytán, M.O., Monjarás Feria, J., Soto, E., Espinosa, N., Benítez, J.M., Georgellis, D., and González-Pedrajo, B. (2018). Novel insights into the mechanism of SepL-mediated control of effector secretion in enteropathogenic *Escherichia coli*. MicrobiologyOpen 7, e00571.

Gendrin, C., Contreras-Martel, C., Bouillot, S., Elsen, S., Lemaire, D., Skoufias, D.A., Huber, P., Attree, I., and Dessen, A. (2012). Structural basis of cytotoxicity mediated by the type III secretion toxin ExoU from *Pseudomonas aeruginosa*. PLoS Pathog. 8, e1002637.

Goure, J., Pastor, A., Faudry, E., Chabert, J., Dessen, A., and Attree, I. (2004). The V antigen of *Pseudomonas aeruginosa* is required for assembly of the functional PopB/PopD translocation pore in host cell membranes. Infect. Immun. 72, 4741–4750.

Halder, P.K., Roy, C., and Datta, S. (2018). Structural and functional characterization of type three secretion system ATPase PscN and its regulator PscL from *Pseudomonas aeruginosa*. Proteins Struct. Funct. Bioinforma.

Hara, N., Morimoto, Y.V., Kawamoto, A., Namba, K., and Minamino, T. (2012). Interaction of the Extreme N-Terminal Region of FliH with FlhA Is Required for Efficient Bacterial Flagellar Protein Export. J. Bacteriol. 194, 5353–5360.

Ho, O., Rogne, P., Edgren, T., Wolf-Watz, H., Login, F.H., and Wolf-Watz, M. (2017). Characterization of the Ruler Protein Interaction Interface on the Substrate Specificity Switch Protein in the *Yersinia* Type III Secretion System. J. Biol. Chem. 292, 3299–3311.

Hu, B., Morado, D.R., Margolin, W., Rohde, J.R., Arizmendi, O., Picking, W.L., Picking, W.D., and Liu, J. (2015). Visualization of the type III secretion sorting platform of *Shigella flexneri*. Proc. Natl. Acad. Sci. U. S. A. 112, 1047–1052.

Hueck, C.J. (1998). Type III Protein Secretion Systems in Bacterial Pathogens of Animals and Plants. Microbiol. Mol. Biol. Rev. 62, 379–433.

Ibuki, T., Imada, K., Minamino, T., Kato, T., Miyata, T., and Namba, K. (2011). Common architecture of the flagellar type III protein export apparatus and F-and V-type ATPases. Nat. Struct. Mol. Biol. 18, 277–282.

Ibuki, T., Uchida, Y., Hironaka, Y., Namba, K., Imada, K., and Minamino, T. (2013). Interaction between FliJ and FlhA, components of the bacterial flagellar type III export apparatus. J. Bacteriol. 195, 466–473.

Imada, K., Minamino, T., Tahara, A., and Namba, K. (2007). Structural similarity between the flagellar type III ATPase FliI and F1-ATPase subunits. Proc. Natl. Acad. Sci. 104, 485–490.

Imada, K., Minamino, T., Uchida, Y., Kinoshita, M., and Namba, K. (2016). Insight into the flagella type III export revealed by the complex structure of the type III ATPase and its regulator. Proc. Natl. Acad. Sci. U. S. A. 113, 3633–3638.

Inoue, Y., Morimoto, Y.V., Namba, K., and Minamino, T. (2018). Novel insights into the mechanism of well-ordered assembly of bacterial flagellar proteins in Salmonella. Sci. Rep. 8, 1787.

Jaquinod, M., Trauchessec, M., Huillet, C., Louwagie, M., Lebert, D., Picard, G., Adrait, A., Dupuis, A., Garin, J., Brun, V., et al. (2012). Mass spectrometry-based absolute protein quantification: PSAQ^TM^ strategy makes use of “noncanonical” proteotypic peptides. PROTEOMICS 12, 1217–1221.

Job, V., Matteï, P.-J., Lemaire, D., Attree, I., and Dessen, A. (2010). Structural basis of chaperone recognition of type III secretion system minor translocator proteins. J. Biol. Chem. 285, 23224–23232.

Journet, L., Agrain, C., Broz, P., and Cornelis, G.R. (2003). The needle length of bacterial injectisomes is determined by a molecular ruler. Science 302, 1757–1760.

Kinoshita, M., Aizawa, S., Inoue, Y., Namba, K., and Minamino, T. (2017). The role of intrinsically disordered C-terminal region of FliK in substrate specificity switching of the bacterial flagellar type III export apparatus. Mol. Microbiol. 105, 572–588.

Kinoshita, M., Tanaka, S., Inoue, Y., Namba, K., Aizawa, S.-I., and Minamino, T. (2020). The flexible linker of the secreted FliK ruler is required for export switching of the flagellar protein export apparatus. Sci. Rep. 10, 838.

Lara-Tejero, M., Kato, J., Wagner, S., Liu, X., and Galán, J.E. (2011). A sorting platform determines the order of protein secretion in bacterial type III systems. Science 331, 1188–1191.

Lee, P.-C., and Rietsch, A. (2015). Fueling type III secretion. Trends Microbiol. 23, 296–300.

Lee, P.-C., Zmina, S.E., Stopford, C.M., Toska, J., and Rietsch, A. (2014). Control of type III secretion activity and substrate specificity by the cytoplasmic regulator PcrG. Proc. Natl. Acad. Sci. 201402658.

Lorenz, C., and Buttner, D. (2009). Functional Characterization of the Type III Secretion ATPase HrcN from the Plant Pathogen *Xanthomonas campestris* pv. vesicatoria. J. Bacteriol. 191, 1414–1428.

Majewski, D.D., Worrall, L.J., Hong, C., Atkinson, C.E., Vuckovic, M., Watanabe, N., Yu, Z., and Strynadka, N.C.J. (2019). Cryo-EM structure of the homohexameric T3SS ATPase-central stalk complex reveals rotary ATPase-like asymmetry. Nat. Commun. 10, 626.

Maresso, A.W., Frank, D.W., and Barbieri, J.T. (2006). CHAPTER 14 - *Pseudomonas aeruginosa* toxins. In The Comprehensive Sourcebook of Bacterial Protein Toxins (Third Edition), J.E. Alouf, and M.R. Popoff, eds. (London: Academic Press), pp. 257–269.

Martinez-Argudo, I., and Blocker, A.J. (2010). The *Shigella* T3SS needle transmits a signal for MxiC release, which controls secretion of effectors. Mol. Microbiol. 78, 1365–1378.

Matson, J.S., and Nilles, M.L. (2001). LcrG-LcrV interaction is required for control of Yops secretion in *Yersinia pestis*. J. Bacteriol. 183, 5082–5091.

Miller, J.H. (1992). A short course in bacterial genetics: a laboratory manual and handbook for Escherichia coli and related bacteria (Plainview, N.Y: Cold Spring Harbor Laboratory Press).

Minamino, T. (2018). Hierarchical protein export mechanism of the bacterial flagellar type III protein export apparatus. FEMS Microbiol. Lett. 365.

Minamino, T., Gonzalez-Pedrajo, B., Yamaguchi, K., Aizawa, S.-I., and Macnab, R.M. (1999). FliK, the protein responsible for flagellar hook length control in Salmonella, is exported during hook assembly. Mol. Microbiol. 34, 295–304.

Minamino, T., Morimoto, Y.V., Hara, N., and Namba, K. (2011). An energy transduction mechanism used in bacterial flagellar type III protein export. Nat. Commun. 2, 475.

Monjarás Feria, J.V., Lefebre, M.D., Stierhof, Y.-D., Galán, J.E., and Wagner, S. (2015). Role of autocleavage in the function of a type III secretion specificity switch protein in *Salmonella enterica* serovar Typhimurium. MBio 6, e01459–01415.

Nickels, B.E. (2009). Genetic assays to define and characterize protein–protein interactions involved in gene regulation. Methods 47, 53–62.

Nilles, M.L., Williams, A.W., Skrzypek, E., and Straley, S.C. (1997). *Yersinia pestis* LcrV forms a stable complex with LcrG and may have a secretion-related regulatory role in the low-Ca2+ response. J. Bacteriol. 179, 1307–1316.

Ple, S., Job, V., Dessen, A., and Attree, I. (2010). Cochaperone Interactions in Export of the Type III Needle Component PscF of *Pseudomonas aeruginosa*. J. Bacteriol. 192, 3801–3808.

Quinaud, M., Chabert, J., Faudry, E., Neumann, E., Lemaire, D., Pastor, A., Elsen, S., Dessen, A., and Attree, I. (2005). The PscE-PscF-PscG complex controls type III secretion needle biogenesis in Pseudomonas aeruginosa. J. Biol. Chem. 280, 36293–36300.

Quinaud, M., Plé, S., Job, V., Contreras-Martel, C., Simorre, J.-P., Attree, I., and Dessen, A. (2007). Structure of the heterotrimeric complex that regulates type III secretion needle formation. Proc. Natl. Acad. Sci. 104, 7803–7808.

Roehrich, A.D., Bordignon, E., Mode, S., Shen, D.-K., Liu, X., Pain, M., Murillo, I., Martinez-Argudo, I., Sessions, R.B., and Blocker, A.J. (2017). Steps for *Shigella* Gatekeeper Protein MxiC Function in Hierarchical Type III Secretion Regulation. J. Biol. Chem. 292, 1705–1723.

Schoehn, G., Di Guilmi, A.M., Lemaire, D., Attree, I., Weissenhorn, W., and Dessen, A. (2003). Oligomerization of type III secretion proteins PopB and PopD precedes pore formation in *Pseudomonas*. EMBO J. 22, 4957–4967.

Schubot, F.D., Jackson, M.W., Penrose, K.J., Cherry, S., Tropea, J.E., Plano, G.V., and Waugh, D.S. (2005). Three-dimensional Structure of a Macromolecular Assembly that Regulates Type III Secretion in Yersinia pestis. J. Mol. Biol. 346, 1147–1161.

Shaulov, L., Gershberg, J., Deng, W., Finlay, B.B., and Sal-Man, N. (2017). The Ruler Protein EscP of the Enteropathogenic *Escherichia coli* Type III Secretion System Is Involved in Calcium Sensing and Secretion Hierarchy Regulation by Interacting with the Gatekeeper Protein SepL. MBio 8.

Shen, D., Quenee, L., Bonnet, M., Kuhn, L., Derouazi, M., Lamotte, D., Toussaint, B., and Polack, B. (2008). Orf1/SpcS chaperones ExoS for type three secretion by *Pseudomonas aeruginosa*. Biomed. Environ. Sci. 21, 103.

Shen, D.-K., Moriya, N., Martinez-Argudo, I., and Blocker, A.J. (2012). Needle length control and the secretion substrate specificity switch are only loosely coupled in the type III secretion apparatus of *Shigella.* Microbiol. Read. Engl. 158, 1884–1896.

Sorg, I., Wagner, S., Amstutz, M., Müller, S.A., Broz, P., Lussi, Y., Engel, A., and Cornelis, G.R. (2007). YscU recognizes translocators as export substrates of the *Yersinia* injectisome. EMBO J. 26, 3015–3024.

Sundin, C., Thelaus, J., Bröms, J.E., and Forsberg, A. (2004). Polarisation of type III translocation by *Pseudomonas aeruginosa* requires PcrG, PcrV and PopN. Microb. Pathog. 37, 313–322.

Wager, B., Faudry, E., Wills, T., Attree, I., and Delcour, A.H. (2013). Current fluctuation analysis of the PopB and PopD translocon components of the *Pseudomonas aeruginosa* type III secretion system. Biophys. J. 104, 1445–1455.

Wagner, S., Stenta, M., Metzger, L.C., Dal Peraro, M., and Cornelis, G.R. (2010). Length control of the injectisome needle requires only one molecule of Yop secretion protein P (YscP). Proc. Natl. Acad. Sci. U. S. A. 107, 13860–13865.

Wagner, S., Grin, I., Malmsheimer, S., Singh, N., Torres-Vargas, C.E., and Westerhausen, S. (2018). Bacterial type III secretion systems: A complex device for delivery of bacterial effector proteins into eukaryotic host cells. FEMS Microbiol. Lett.

Wood, S.E., Jin, J., and Lloyd, S.A. (2008). YscP and YscU switch the substrate specificity of the *Yersinia* type III secretion system by regulating export of the inner rod protein YscI. J. Bacteriol. 190, 4252–4262.

Yahr, T.L., Barbieri, J.T., and Frank, D.W. (1996). Genetic relationship between the 53- and 49-kilodalton forms of exoenzyme S from *Pseudomonas aeruginosa*. J. Bacteriol. 178, 1412–1419.

Yang, H., Shan, Z., Kim, J., Wu, W., Lian, W., Zeng, L., Xing, L., and Jin, S. (2007). Regulatory Role of PopN and Its Interacting Partners in Type III Secretion of *Pseudomonas aeruginosa*. J. Bacteriol. 189, 2599–2609.

Yoshida, Y., Miki, T., Ono, S., Haneda, T., Ito, M., and Okada, N. (2014). Functional characterization of the type III secretion ATPase SsaN encoded by *Salmonella* pathogenicity island 2. PloS One 9, e94347.

Yu, X.-J., Grabe, G.J., Liu, M., Mota, L.J., and Holden, D.W. (2018). SsaV Interacts with SsaL to Control the Translocon-to-Effector Switch in the Salmonella SPI-2 Type Three Secretion System. 9, 16.

Zarivach, R., Vuckovic, M., Deng, W., Finlay, B.B., and Strynadka, N.C.J. (2007). Structural analysis of a prototypical ATPase from the type III secretion system. Nat. Struct. 38 Mol. Biol. 14, 131–137.

