## Supplementary Material for "The PopN gate-keeper complex acts on the ATPase PscN to regulate the T3SS secretion switch from early to middle substrates in *Pseudomonas aeruginosa*"

**Table S1: Primers and DNA fragment used in this work.**

| Name | Sequence 5’- 3’ |
|---|---|
| Delta_pscN_1 | CCG GGC GCC TCG AGC TTC TGC TG |
| Delta_pscN_2 | ATG GCG CTG ATC CAG CGC CTG GTG CTG CTG |
| Delta_pscN_3 | TGG ATC AGC GCC ATG CGG CC |
| Delta_pscN_4 | CCG GGA GCG GAG CCG TAT CCA C |
| NdeI_**Strep**_PscN | AAC CAT ATG **GCC AGC TGG AGC CAC CCG CAG TTC GAG AAG CCG GGC** ATG CCC<br>GCG CCT CTC TCT CCT C |
| PscN_Hind | TGG AAG CTT TCA TGC CGA GAG GCT CCG CAA CTG CGC G |
| NcoI_**FSS**_NterPscN_Not | CCATGGGCTCGAGCTCC**GACTACAAGGACGACGACGACAAG**TCGAGCTCCGGCAGCGC<br>GAGCGCG**TGGAGCCACCCGCAGTTCGAGAAG**GGCGGCGGCAGCGGCGGCGGCAGCGG<br>CGGCAGCGCG**TGGAGCCACCCGCAGTTCGAGAAG**ATCGAGGGAAGGCATATGCCCGCG<br>CCTCTCTCTCCTCTCATCGTCCGGATGCGCCACGCCATCGAAGGCTGCCGGCCGATCCAGA<br>TCCGCGGGCGGGTCACCCAGGTCACCGGAACCCTGCTCAAGGCCGTGGTGCCCGGCGTGC<br>GCATCGGCGAACTCTGCCAGTTGCGCAATCCCGACCAGAGCCTGGCGCTGCTCGCCGAGG<br>TCATCGGCTTCCAGCAGCACCAGGCGCTGCTCACCCCGCTCGGCGAGATGCTCGGGGTTT<br>CCTCCAACACCGAAGTCAGCCCTACCGGCGGCATGCATCGCGTGGCGGTCGGAGAGCACC<br>TGCTCGGGCAGGTGCTCGACGGTCTCGGCCGCCCCTTCGACGGCAGCCCGCCGGCCGAGC<br>CGGCGGCCTGGTATCCGGTCTACCGGGATGCCCCGCAACCGATGAGCCGGCGCCTGATAG<br>AGCGGCCGCT |
| PopNEX5 | AAA AAC ATA TGG ACA TCC TCC AGA GTT CCT |
| Pcr2fB-3 | AAT CCG CTC AAC AGA TGA TTC ATG CGG CGA GCA CTT CGC T |
| PscBf2-5 | TCG GCG CGC AGC GAA GTG CTC GCC GCA TGA ATC ATC TGT TGA GCG GAT T |
| PscB3H | AAA AAA AGC TTT CAG GGA CGC CAC ACC GGA GCC T |
| PopN-5NotI | AAA AAG CGG CCG CAG ACA TCC TCC AGA GTT CCT CC |
| PopN-3AscI | AAA AAG GCG CGC CTC AGA AGG CCC GTA TGC CAT |
| PscB-5NotI | AAA AAG CGG CCG CAG ATC ATC TGT TGA GCG GAT TG |
| PscB-3AscI | AAA AAG GCG CGC CTC AGG GAC GCC ACA CCG GAG CCT G |
| Pcr1-5NotI | AAA AAG CGG CCG CAG CAT ACG GGC CTT CTG AAT TGA C |
| Pcr1-3AscI | AAA AAG GCG CGC CTC AAC CCA GTC CAT GCT GCT GCT C |
| Pcr2-5NotI | AAA AAG CGG CCG CAG ACT GGG TTG AGC TGG CCG TC |
| Pcr2-3AscI | AAA AAG GCG CGC CTC ATG CGG CGA GCA CTT CGC TG |
| PscBf2-5 | TCG GCG CGC AGC GAA GTG CTC GCC GCA TGA ATC ATC TGT TGA GCG GAT T |
| Pcr2fB-3 | AAT CCG CTC AAC AGA TGA TTC ATG CGG CGA GCA CTT CGC T |

**Table S2: Affinity between PscN and T3SS secreted proteins**. The $K_D$s (nM) between exported cargos, their chaperones alone or in complex and the ATPase PscN were measured by MST. ND: non-determined

<table>
<tr><th colspan="4" rowspan="2">Protein</th><th>K<sub>D</sub> (nM)</th><th>K<sub>D</sub> (nM)</th><th>K<sub>D</sub> (nM)</th></tr>
<tr><th>Two-partners buffer*</th><th>In the absence of PopN complex</th><th>In the presence of PopN complex</th></tr>
<tr><td colspan="2" rowspan="3">Effector</td><td>Complex</td><td>ExoU-SpcU</td><td>45 ± 13</td><td>47 ± 17</td><td>46 ± 13</td></tr>
<tr><td>Cargo</td><td>ExoU</td><td>32 ±12</td><td>ND</td><td>ND</td></tr>
<tr><td>Chaperone</td><td>SpcU</td><td>41722 ±15413</td><td>ND</td><td>ND</td></tr>
<tr><td rowspan="7">Translocator</td><td rowspan="4">Translocon</td><td>Complex</td><td>PcrH-PopB</td><td>4939 ±170</td><td>6722 ± 477</td><td>12365 ± 1361</td></tr>
<tr><td>Complex</td><td>PcrH-PopD</td><td>4719 ±1772</td><td>ND</td><td>ND</td></tr>
<tr><td>Cargo</td><td>PopB/PopD</td><td>ND</td><td>ND</td><td>ND</td></tr>
<tr><td>Chaperone</td><td>PcrH</td><td>31597 ± 6245</td><td>ND</td><td>ND</td></tr>
<tr><td rowspan="3">Tip</td><td>Complex</td><td>PcrV-PcrG</td><td>ND</td><td>ND</td><td>ND</td></tr>
<tr><td>Cargo</td><td>PcrV</td><td>2648 ± 220</td><td>4205 ± 623</td><td>13104 ± 2475</td></tr>
<tr><td>Chaperone</td><td>PcrG</td><td>ND</td><td>ND</td><td>ND</td></tr>
<tr><td colspan="2" rowspan="3">Needle</td><td>Complex</td><td>PscE-PscF-PscG</td><td>>150000</td><td>>45000</td><td>5607 ± 2206</td></tr>
<tr><td>Cargo</td><td>PscF</td><td>ND</td><td>ND</td><td>ND</td></tr>
<tr><td>Chaperone</td><td>PscE-PscG</td><td>>150000</td><td>ND</td><td>ND</td></tr>
<tr><td colspan="2" rowspan="3">Gate-keeper</td><td>Complex</td><td>PopN-Pcr1-Pcr2-PscB</td><td>806 ± 105</td><td>1174 ± 40</td><td>ND</td></tr>
<tr><td>Cargo</td><td>PopN-Pcr1</td><td>ND</td><td>ND</td><td>ND</td></tr>
<tr><td>Chaperone</td><td>Pcr2-PscB</td><td>ND</td><td>ND</td><td>ND</td></tr>
</table>

* The final buffer composition of this set of assays is somehow different from the one for the assays in the presence or absence of the PopN complex.

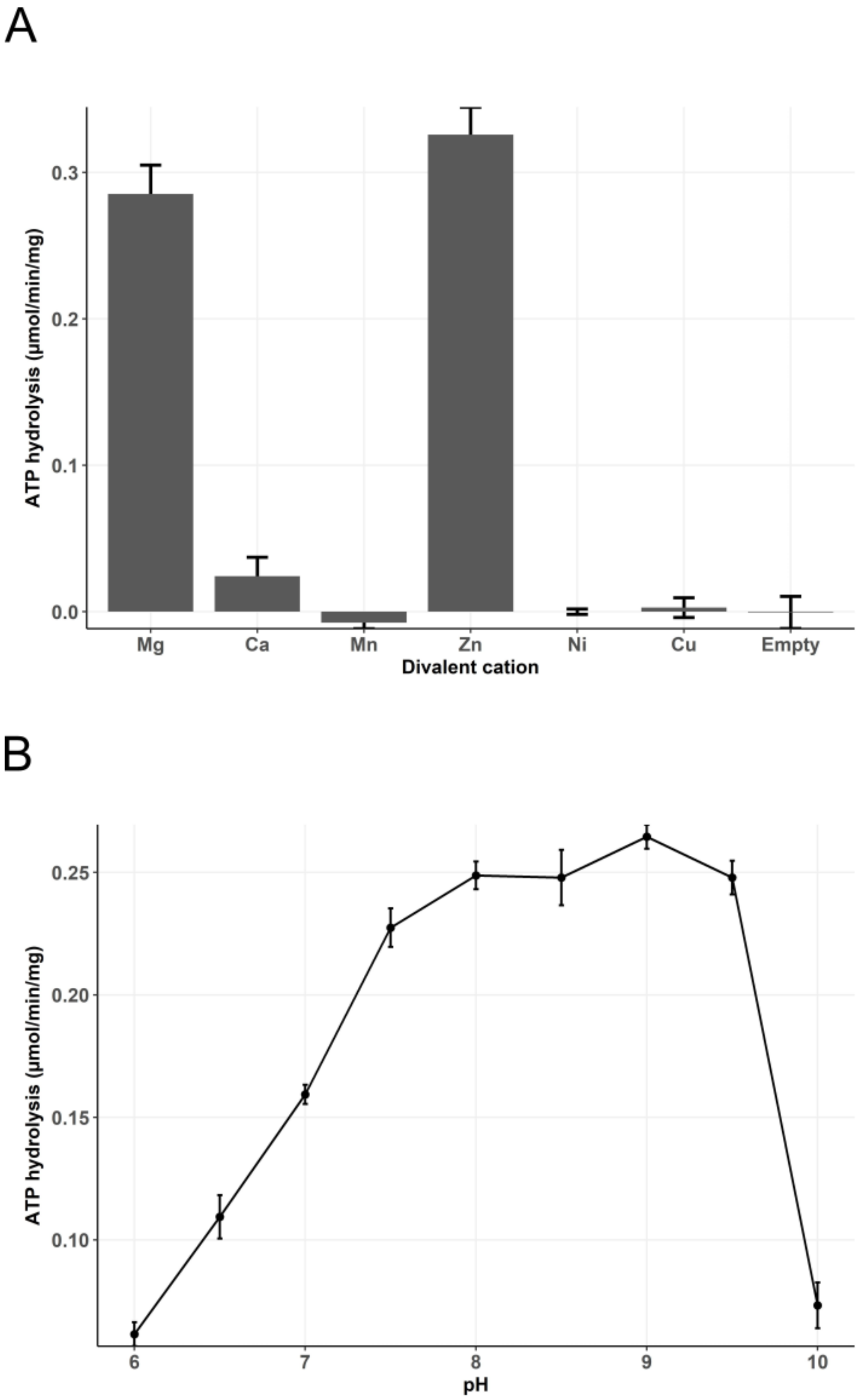


**Figure S1. ATPase activity of PscN.** Purified FSS-PscN was incubated with 5 mM ATP and 5 mM of the indicated divalent cations at pH 9 (A) or with 5 mM ATP and 5 mM $MgCl_2$ at the indicated pHs (B) for 30 minutes. The released phosphate was measured using the malachite green method.

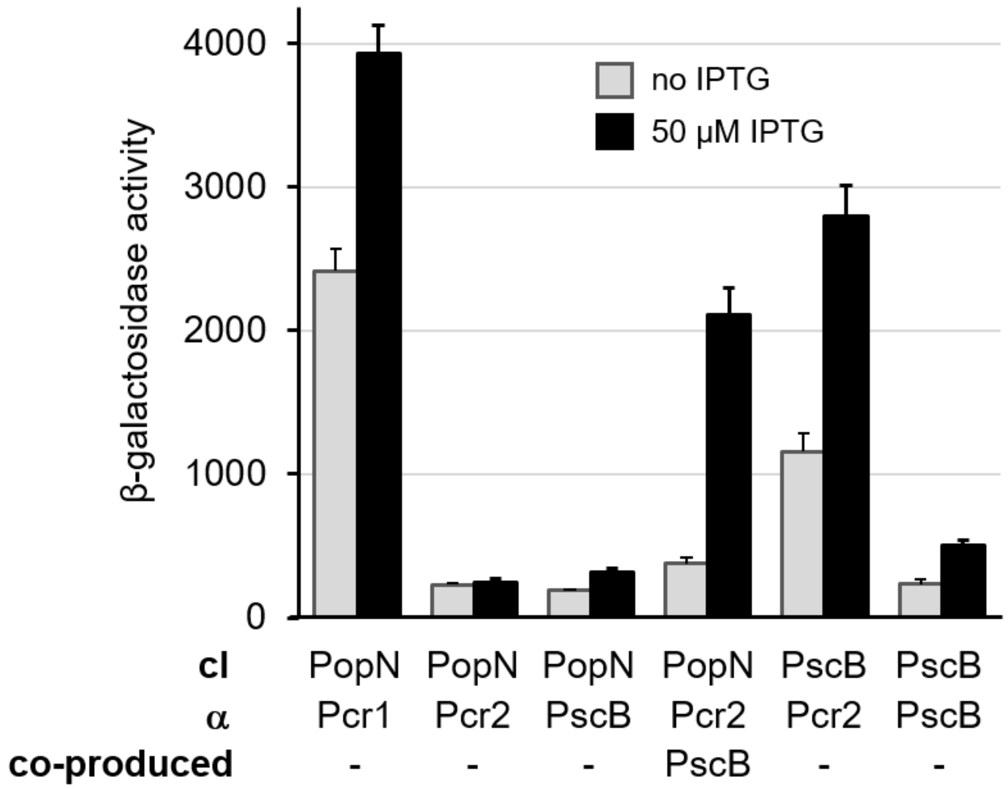


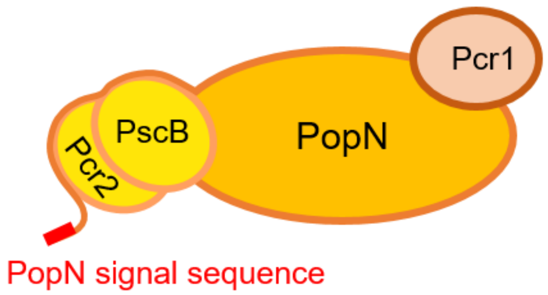


**Figure S2. Two-hybrid characterization of the PopN complex.**

Pairwise interactions among PopN-complex proteins were examined using the alpha-cI bacterial two-hybrid system. Fusion partners to RNA-polymerase alpha, or lambda cI are indicated below the graph and were co-produced (IPTG induction) in an *E. coli* reporter strain. Interaction between the cI and alpha fusion proteins recruits RNA-polymerase to a test promoter activating *lacZ* transcription, which was assayed by ß-galactosidase assay. To demonstrate the interaction between PopN and the hetero-dimeric Pcr2/PscB chaperone, *pscB* was co-expressed together with the alpha-*pcr2* fusion by translationally coupling the *pscB* open reading frame to the alpha-*pcr2* gene. A cartoon of the PopN complex is shown below the graph. The organization is based on the organization of the homologous YopN regulatory complex from *Yersinia pestis* (Schubot et al., 2005).

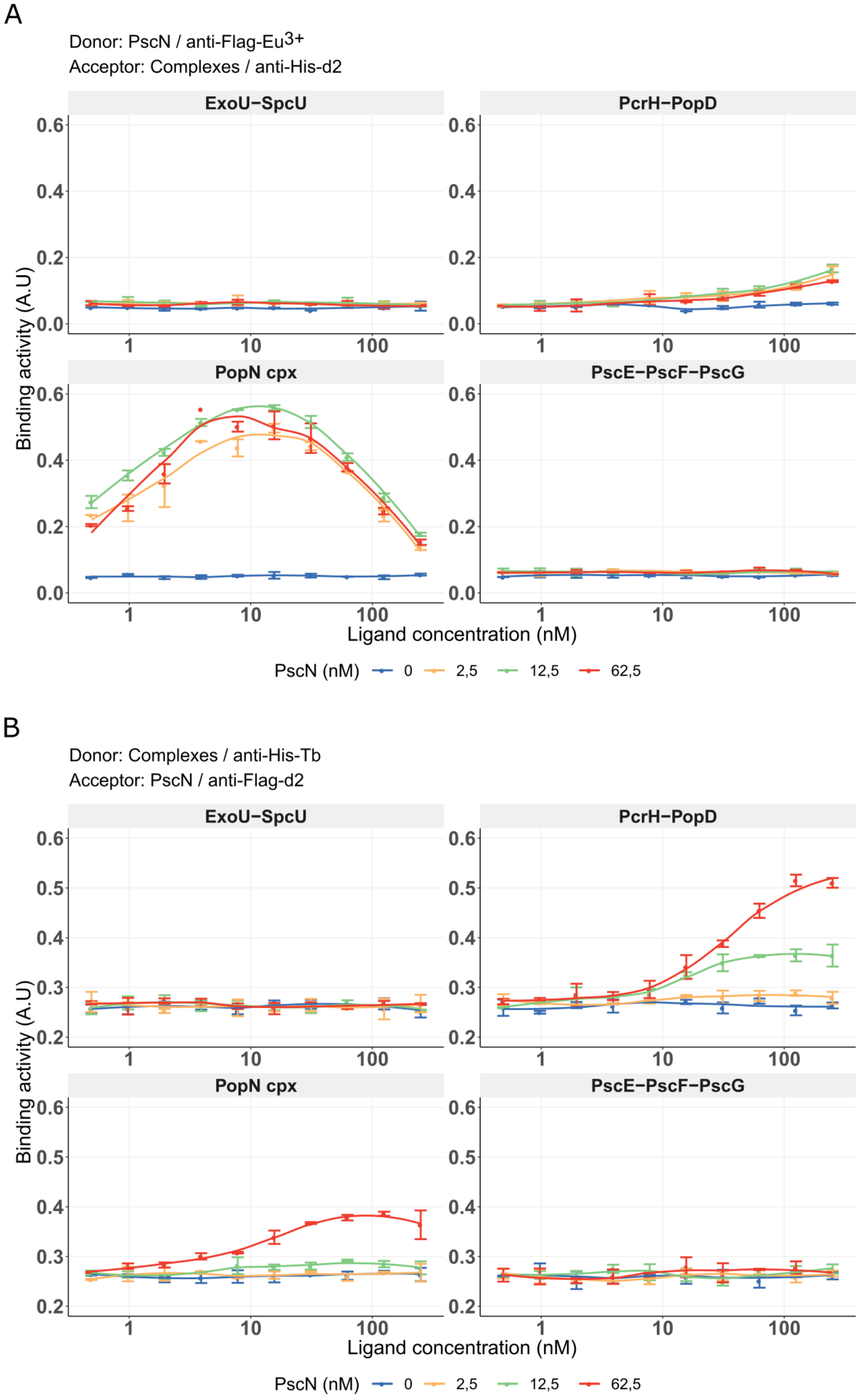


**Figure S3: HTRF detection of the binding of the ATPase PscN to the exported protein complexes.** Effector (ExoU-SpcU), translocator (PcrH-PopD), needle (PscE-PscF-PscG) or gate-keeper (PopN cpx) complexes expressed as His-tagged fusion proteins were mixed with Flag-tagged PscN at the indicated concentrations in the presence of two pairs of fluorophore-tagged antibodies anti-Flag-$Eu^{3+}$ and anti-His-d2 (A) or anti-Flag-d2 and anti-His-Tb (B). The binding activity was detected by measuring time-resolved fluorescence transfer.

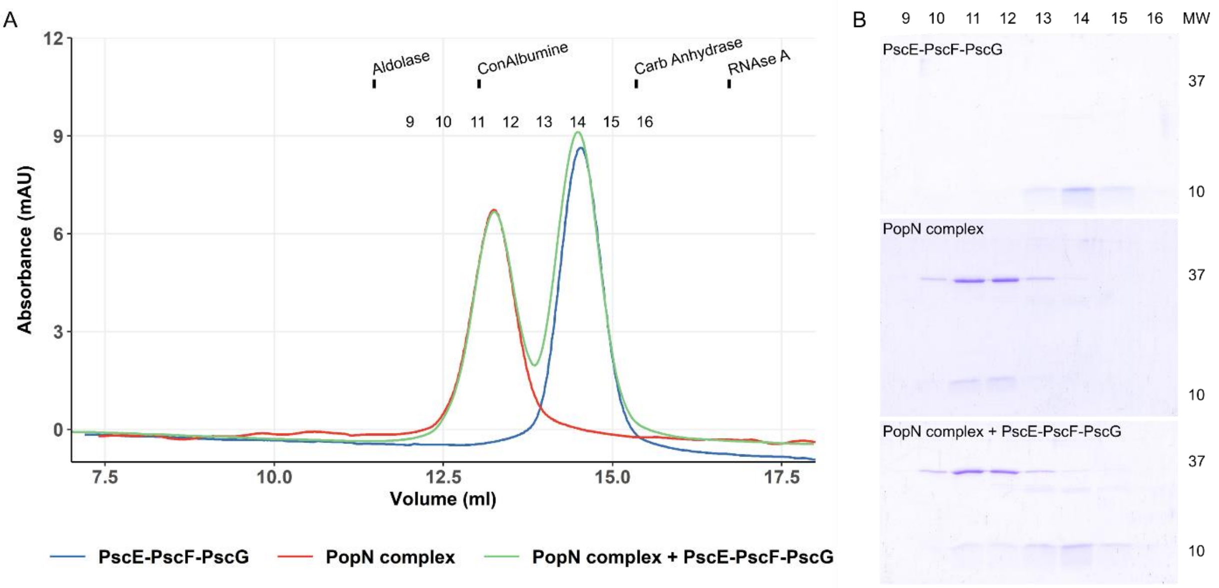


**Figure S4: Size exclusion chromatography analysis of PopN complex and PscF complex.** PscF complex at 5 µM, PopN complex at 1 µM or a mixture of the two complexes were injected onto a Superdex 200 increase column. A) The two peaks obtained in the mixture are similar to each peak from the two complexes alone, indicating an absence of interaction between PscF and PopN complexes. Elution volumes of Aldolase (158 kDa), ConAlbumine (75 kDa), Carbonic Anhydrase (29 kDa) and RNAse A (13.7 kDa) are indicated by dashes and the numbers indicate the collected fractions. B) Fractions of each chromatography were analyzed by 18% SDS-PAGE showing that the fraction content the two complexes eluted separately when they were injected together on the column. In this electrophoresis, the small Pcr1, Pcr2, PscB, PscE, PscF and PscG could not be resolved despite many attempts and migrate as smears. Theoretical molecular masses: His-PopN 33.7 kDa; Pcr1 10.4 kDa; Pcr2 13.7 kDa; PscB 15.4 kDa; PscE 9.5 kDa; PscF 9.3 kDa and PscG 12.5 kDa

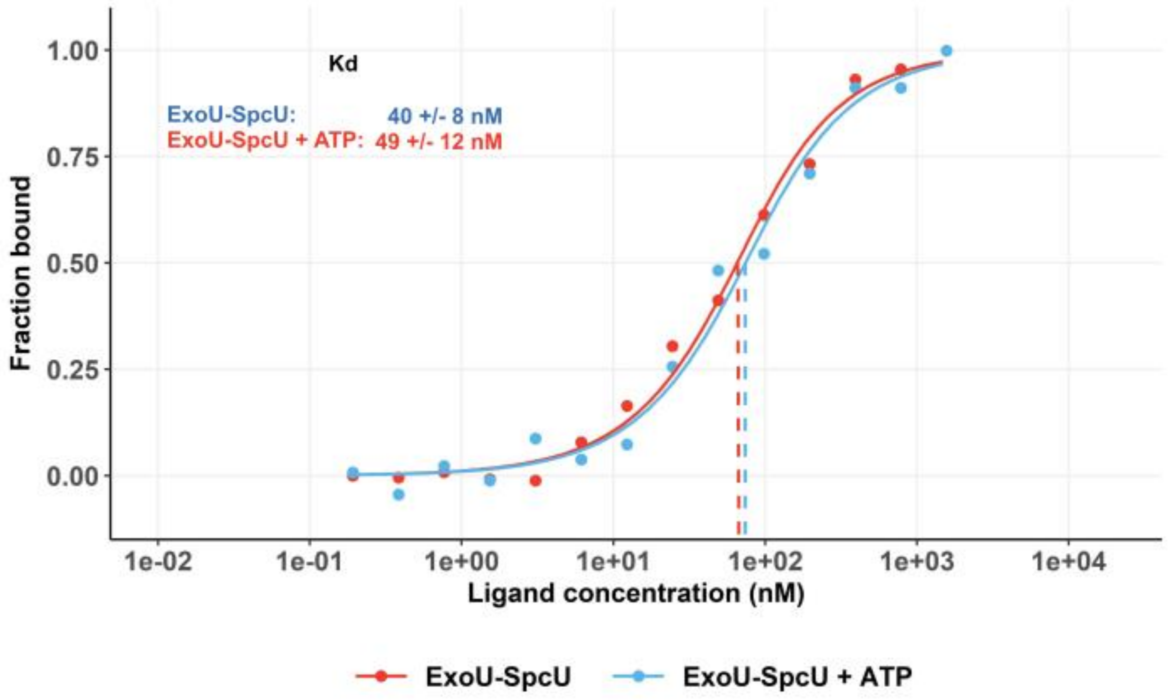


**Figure S5: The presence of ATP does not alter the affinity between PscN and ExoU-SpcU.** FSS-PscN was labeled with MST dye and 100 nM of labeled protein were incubated with different concentrations of ExoU-PscU in the absence or presence of 5 mM ATP. A titration fitted curve for each ligand, which represents the bound fraction depending on ligand concentrations, and $K_D$ were obtained.

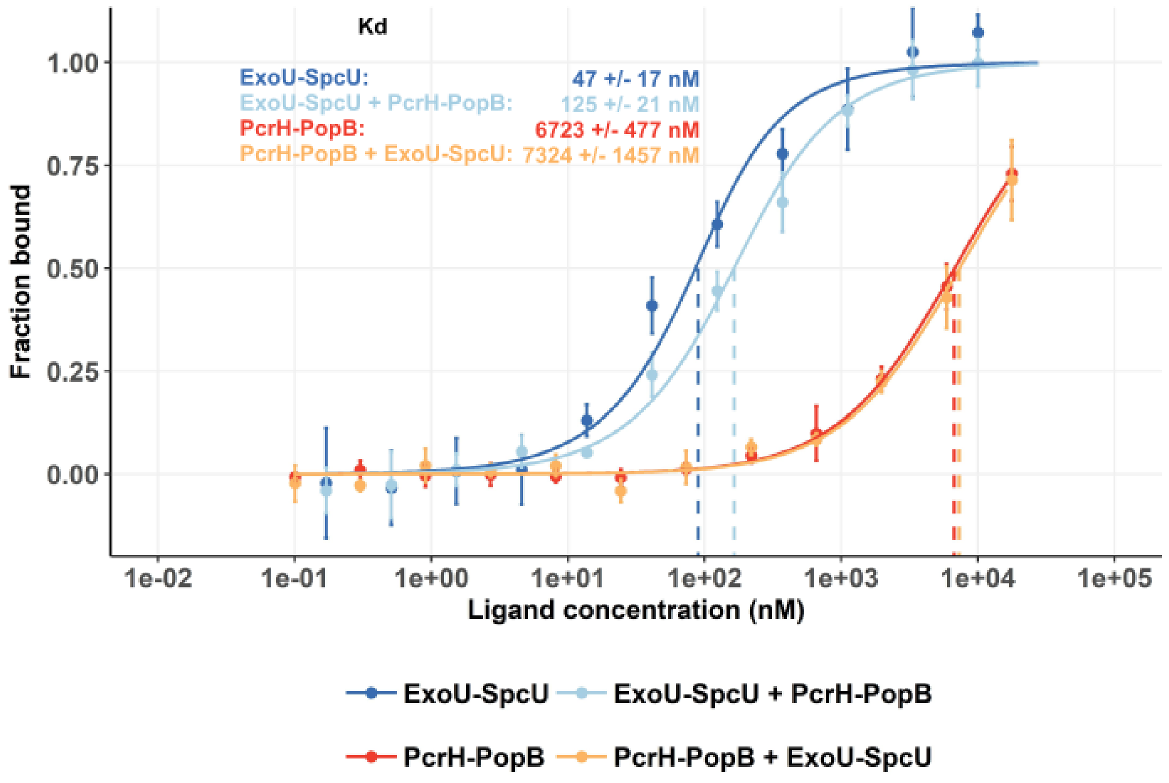


**Figure S6: Competition between secreted proteins for the interaction with PscN.** The presence of PcrH-PopB slightly decreases the affinity between ExoU-SpcU and PscN while the binding of the effector complex does not affect the affinity between the ATPase and the translocator complex.
